# A conserved protein inhibitor brings under check the activity of RNase E in cyanobacteria

**DOI:** 10.1101/2023.03.20.533394

**Authors:** Su-Juan Liu, Gui-Ming Lin, Yu-Qi Yuan, Wenli Chen, Ju-Yuan Zhang, Cheng-Cai Zhang

## Abstract

RNase E is a major ribonuclease for RNA metabolism in bacteria. Because it has a large substrate spectrum and poor substrate specificity, its activity must be well controlled under different conditions. Only a few regulators of RNase E are known in bacteria, limiting our understanding on the posttranscriptional regulatory mechanisms operating in these organisms. Here we show that, RebA, a protein universally present in cyanobacteria, interacts with RNase E in the filamentous cyanobacterium *Anabaena* PCC 7120. Distinct from those known regulators of RNase E, RebA interacts with the 5’ sensor domain in the catalytic region of RNase E, and suppresses the cleavage activities of RNase E for all tested RNA substrates irrespective of their 5’-end status. Consistent with the inhibitory function of RebA on RNase E, conditional depletion of RNase E and overproduction of RebA caused formation of elongated cells, whereas the absence of RebA and overproduction of RNase E resulted in a shorter-cell phenotype. We further showed that the morphological changes caused by altered levels of RNase E or RebA are dependent on their physical interaction. The action of RebA represents a new mechanism, highly conserved in cyanobacteria, for RNase E regulation. Our findings provide insights into the regulation and the function of RNase E, and demonstrate the importance of balanced RNA metabolism in bacteria.

## INTRODUCTION

RNA degradation is one of the essential processes for RNA metabolism in all organisms. In bacteria, mRNAs usually have an average lifetime of only a few minutes (Bernstein et al., 2002; Hambraeus *et al*., 2003; Andersson *et al*., 2006; Steglich *et al*., 2010). The rapid degradation of mRNA molecules serves as an important mechanism for rapid adjustment of protein levels in response to the changing environmental conditions. In *E. coli* and many other bacteria, the endoribonuclease RNase E plays a central role in mRNA degradation; it also participates in the maturation of ribosomal RNAs, tRNAs, as well as many small regulatory RNAs (Mackie, 2013; Zhang et al., 2022). RNase E homologs from most bacteria share a very similar overall architecture, with a conserved catalytic N-terminal half and a divergent noncatalytic C-terminal half (Aït-Bara and Carpousis, 2015). RNase E from *E. coli* functions as a tetramer. Its N-terminal half (1–529 aa) includes several subdomains: RNase H-like domain, S1 domain that contributes to substrate binding, the 5′ monophosphate sensing domain (5’ sensor domain) that binds to the 5’ p of monophosphorylated RNA substrates, DNase I domain that harbors the RNA hydrolyzing center, Zn-link and a small folded domain that are involved in tetramerization. The C-terminal half (530-1061 aa), though largely disordered, also contains several functional motifs, including two RNA binding sites, and several protein recognition sites that can recruit the exoribonuclease PNPase, the RNA helicase RhlB, and the glycolytic enzyme enolase to form the RNA degradation complex called RNA degradosome (Py et al., 1996; Miczak et al., 1996).

RNase E cuts single-stranded RNA substrates preferentially at AT-rich sites with only modest sequence specificity (McDowall et al., 1994; McDowall et al., 1995; Huang et al., 1998; Chao et al., 2017; Hoffmann et al., 2021). At least two potential pathways for substrate recognition by RNase E have been identified: a 5′ end-dependent mode that relies on 5′ monophosphate recognition (Callaghan et al., 2005; Garrey et al., 2009; Bandyra et al., 2018), and an internal entry mode by which substrates are cleaved regardless of their 5′ end phosphorylation status (Kime et al., 2010; Clarke et al., 2014). In the 5′ end-dependent mode, the 5’ p of the substrate binds to a pocket residing within the 5’ sensor domain of RNase E. Such a binding mode could facilitate orientation of the substrates to the reaction center and allow RNase E to scan linearly from the 5′ end for sites of cleavage (Callaghan et al., 2005; Garrey et al., 2009; Bandyra et al., 2018; Richards and Belasco, 2019). The mechanism for the direct entry mode is less clear, and may involve the recognition of a duplex region in the substrate by several residues from the RNase H-like domain and the small domain (Bandyra et al., 2018).

The RNase E encoding gene, *rne*, is essential in all bacteria investigated. RNase E depletion in *E. coli* could increase the average half-life of bulk mRNA from about 2.5 min to over 10 min (Ono and Kuwano, 1979; Babitzke and Kushner, 1991). RNase E, together with RNase III, could account for the initiation of decay of ∼72% of the *E. coli* transcriptome (Stead et al., 2011). Since RNase E has such a global impact on RNA metabolism with poor substrate selectivity, its cellular activity needs to be properly controlled. However, only a few mechanisms on the regulation of RNase E activity have been identified in a limited bacterial species. For instances, *E. coli* RNase E can maintain the abundance of its own transcripts to a proper level by cleaving at its own 5’-UTR region (Jain and Belasco, 1995). Additionally, several proteins regulate the activity of RNase E by physical interactions. For example, the ribonuclease activity A (RraA) binds to the two RNA binding sites of the noncatalytic region of RNase E, and interfere with the substrate binding activity of RNase E (Lee *et al*., 2003; Górna et al., 2010). The *E. coli* ribosomal protein L4 could interact with different sites of the C-terminal region of RNase E and inhibit RNase E activity on the artificial substrate LU13 *in vitro*, but it seems to only affect the degradation of certain mRNAs involved in stress-response *in vivo* (Singh et al., 2009). Dip, a protein from the giant phage фKZ, could inhibit the activity of RNase E of its host *Pseudomonas aeruginosa* by binding to the RNA binding sites in the noncatalytic region (Van den Bossche et al., 2016). Recently, a bacterial cell wall peptidoglycan hydrolase, AmiC, was shown to stimulate the activity of RNase E, potentially by enhancing RNase E multimerization (Moore et al., 2021). It is yet unknown if any of these regulation mechanisms exist in the other RNase E-containing organisms.

Cyanobacteria, capable of oxygen-evolving photosynthesis and evolutionarily related to chloroplasts of higher plants, are a unique phylum of gram-negative prokaryotes with great morphological diversity (Stanier and Cohen-Bazire, 1977). An *rne* gene is present in each sequenced cyanobacterial genome, and attempts to inactivate the *rne* gene in several cyanobacterial species all failed (Horie et al., 2007; Cameron et al., 2015; Cavaiuolo et al., 2020), implying that this endoribonuclease has an essential role in cyanobacteria. Cyanobacterial RNase E proteins also consist of a conserved catalytic domain at the N-terminal and an intrinsically disordered non-catalytic region at the C-terminal (Zhang et al., 2014). The catalytic region is similar to that of *E. coli* but lacks the small folded domain, while the noncatalytic region shows no detectable similarity. The non-catalytic region shows no detectable similarity to that of *E. coli* RNase E, but contains several subregions well conserved in cyanobacteria. In the filamentous cyanobacterium *Anabaena* PCC 7120 (*Anabaena* hereafter), RNase E can interact with the exoribonuclease PNPase, the exoribonuclease RNase II, and the RNA helicase CrhB, suggesting the presence of an RNase E-based RNA degradosome in this organism (Zhang et al., 2014; Zhou et al., 2020; Yan et al., 2020).

Cyanobacterial RNase E has a catalytic properties and substrate preference similar to its counterparts from other bacteria *in vitro*, and acts on a large panel of RNA species (Kaberdin et al., 1998; Horie et al., 2007; Zhang et al., 2014; Behler et al., 2018; Hoffmann et al., 2021). The transcriptome-wide cleavage sites of RNase E were mapped in the unicellular cyanobacterial *Synechocystis* PCC 6803, revealing a consensus site featured with adenine residues at positions −4 and −3 upstream and uridine residues immediately downstream of the cleavage site, especially at position +2 (Hoffmann et al., 2021). *Synechocystis* RNase E has been shown to participate in regulating various cellular activities, such as photosynthesis, plasmid replication, and the maturation of crRNAs and tRNAs (Horie et al., 2007, Sakurai et al., 2012; Hoffmann et al., 2021). Nevertheless, how the activity of RNA degradation is controlled in cyanobacteria remains little understood. Here, we report the identification of RebA, previously annotated as a protein of unknown function, by co-immunoprecipitation with RNase E from the cell lysate of *Anabaena*. We show that this protein binds to the catalytic domain of RNase E and inhibits its binding and cleavage activity on various RNA substrates, and its action on RNase E plays a role in cell morphology control.

## RESULTS

### Identification of RebA, a novel RNase E-binding protein, universally present in cyanobacteria

Aiming to identify new components of the RNA degradosome or RNase E regulators in *Anabaena*, we performed a co-immunoprecipitation (CoIP) assay using the polyclonal antibodies against RNase E. The proteins precipitated were separated on SDS-PAGE and each gel band was subjected to mass spectrometry analysis (Figure 1A). Most of the bands corresponded to degraded fragments of RNase E, likely due to that the intrinsically disordered nature of the noncatalytic region of RNase E makes the protein very susceptible to proteolytic degradation. Among the known partners of RNase E (i.e., RNase II, PNPase and CrhB), only RNase II was detected in one band; the missing of the others could be due to their lower affinity to RNase E or the partial degradation of their recognition regions in RNase E. Interestingly, there existed one band corresponding to a protein of unknown function (protein id: All1338) together with a degraded product of RNase E. Using bacterial two-hybrid assays, we further showed that All1338 interacted directly with RNase E, but not with the three known RNase E partners, RNase II, PNPase and CrhB (Figure 1B). These results demonstrated that All1338 is a new interacting partner of RNase E. We thus named All1338 as RebA (<u>R</u>Nase <u>E</u> <u>b</u>inding protein <u>A</u>) in this study.

**Figure 1.**
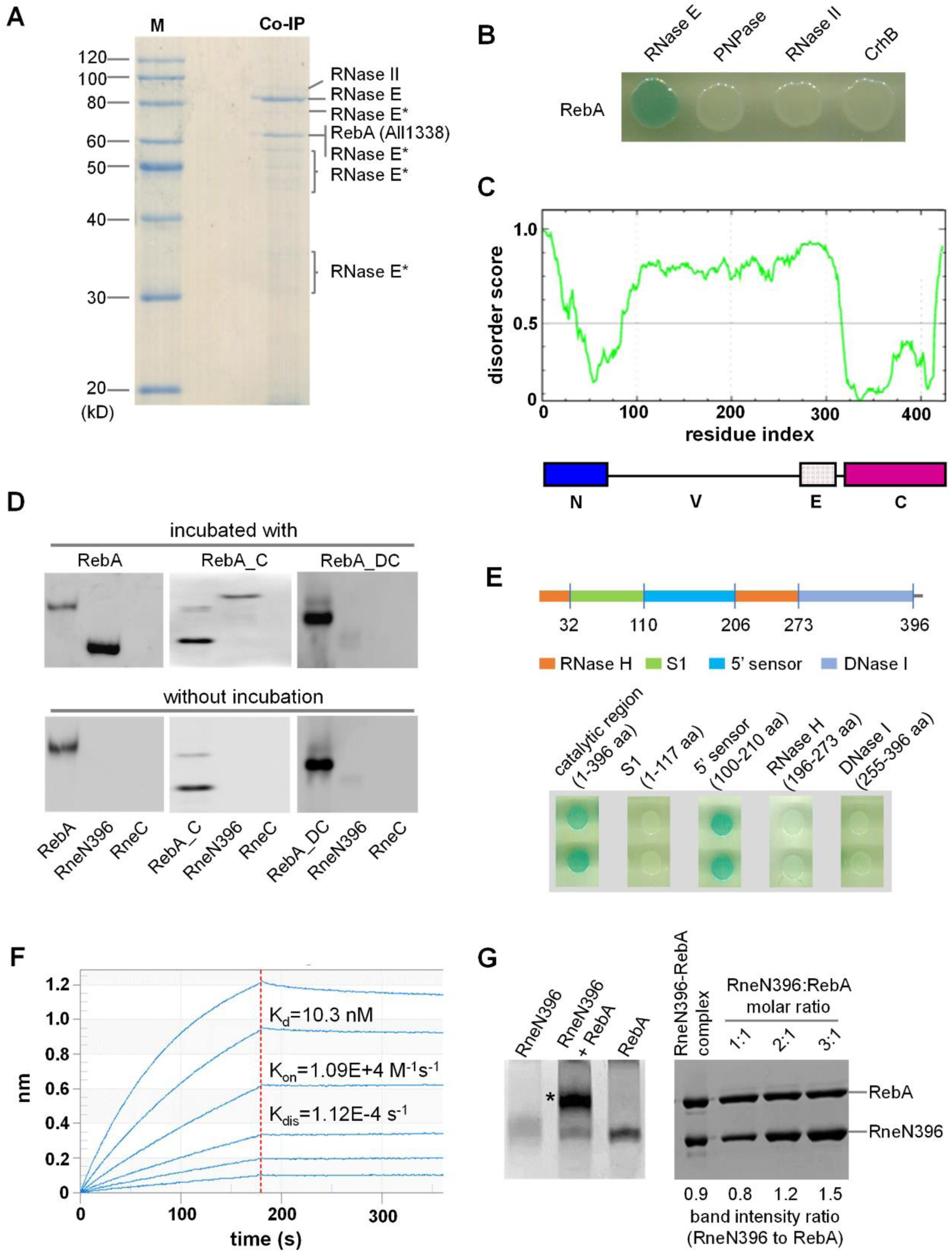
RebA interacts with RNase E both *in vivo* and *in vitro*. **(A)** Identification of the proteins in complex with RNase E by co-immunoprecipitation from *Anabaena* cell lysate, using polyclonal antibodies against *Anabaena* RNase E. The co-immunoprecipitated proteins were separated by a 10% SDS-PAGE, and the proteins in the visible bands after Coomassie-blue staining were identified by mass spectrometry. Those bands corresponding to the degraded products of RNase E were marked as “RNase E*”. **(B)** Bacterial two hybrid assay for the interaction between RebA and RNase E. No interaction was found between RebA and other known RNase E partners. **(C)** Disorderity (upper) and domain (lower) analyses for RebA. Prediction of intrinsically disordered residues in RebA was performed with PONDR-FIT (http://original.disprot.org/pondr-fit.php). The line at 0.5 of Y-axis is the threshold above which it is predicted to be disordered. The domain architecture of RebA was defined based on the alignment of RebA homologs: N, the N-terminal conserved region; V, a highly variable region; E, a region rich in glutamic acid and aspartic acid; C, the C-terminal conserved region. **(D)** Determination of the main regions within RebA and RNase E responsible for interaction by Far-Western blotting assays. The N-terminal region of RNase E (RneN396) and the C-terminal region of RNase E (RneC), together with RebA, RebA_C (C-terminal region of RebA) or RebA_DC (RebA without the C-terminal region), were separated on two sets of SDS-PAGE gels, followed by transfer onto PVDF membranes. One set of the membranes (upper) were incubated respectively with purified RebA, RebA_C or RebA_DC, and detected with the polyclonal antibodies against RebA. The other set of the membranes (bottom) were incubated with BSA prior to the detection with RebA antibodies. **(E)** Interaction of different domains of RNase E (RNase H, S1, 5’ sensor and DNase I, upper panel) with RebA, determined using bacterial two-hybrid assays, with the full-length catalytic region of RNase E as a positive control (lower panel). **(F)** Quantification of RebA-RNase E interaction using surface plasmon resonance (SPR) assay. The association rate (K_on_), dissociation rate (K_dis_), and dissociation constant (K_d_) were estimated from the sensorgram. **(G)** Stoichiometry of RebA-RNase E complex. RneN396, RebA and a mixture of RneN396 and RebA were incubated for 30 min, then separated on a native gel (left gel). The band of the protein complex (indicated by an “*”) was excised and analyzed with SDS-PAGE (right gel), with purified RneN396 and RebA at defined molar ratios loaded for comparison. The intensity of the protein bands in the SDS-PAGE was quantified with Image J.

A RebA homolog could be found in each sequenced cyanobacterial genome by BLAST analysis. The length of RebA homologs varies greatly, ranging from 218 aa in the marine picocyanobacterium *Prochlorococcus* CCMP 1986 to 423 aa in *Anabaena* PCC 7120. The phylogenetic tree of RebA homologs from representative strains has a topology similar to the phylogenetic tree of cyanobacterial species (Figure S1; Shih et al., 2013), indicating that *rebA* is an ancient gene that has already existed in the common ancestor of cyanobacteria. Outside cyanobacteria, RebA homologs are only confidently identified in several species of photosynthetic *Paulinella*, and these *Paulinella* sequences are grouped together with those of marine *Synechococcus* and *Prochlorococcus* strains (Figure S1), suggesting their cyanobacterial origin. Indeed, photosynthetic *Paulinella* is a lineage of amoebae that had obtained the capacity of photosynthesis 90–140 million years ago via endosymbiosis of a cyanobacterium belonging to the *Prochlorococcus*-*Synechococcus* lineage (Gabr et al., 2020).

A Pfam search showed that the major part of RebA did not match to any known protein domains, except that its first 42 residues displayed a low similarity (e-value = 0.00022) to a helix-turn-helix domain. We also performed a disorderity analysis on RebA using PONDR-FIT. It showed that the N-terminal and C-terminal regions of RebA are ordered with high probability, while the middle part is disordered (Figure 1C). Sequence alignment with RebA and its homologs showed that the protein can be divided into four distinct domains, which we named as N, V, E and C, respectively (Figure 1C). The domains N and C are two conserved regions located at the N-terminus (∼60 aa) and C-terminus (∼90 aa), respectively, corresponding to the two ordered regions identified by PONDR-FIT analysis. The highly variable V domain (∼200 aa) and the glutamic acid-rich, negatively charged E domain (∼50 aa), are located between N and C. The different lengths of RebA homologs are mainly due to the length variation of their V and E regions.

### The C domain of RebA interacts with the 5’ sensor domain of RNase E

We next identified the regions in RebA and RNase E responsible for interaction. As the C domain of RebA is the most conserved, we first tested its role in interaction. We expressed and purified, respectively, the recombinant proteins of the full-length RebA, the C region of RebA (RebA_C), the truncated form of RebA without the C region (RebA_DC), the catalytic core of RNase E (RneN396, corresponding to 1-396 aa) and the noncatalytic domain of RNase E (RneC, corresponding to 401-687 aa). RebA, RebA_C and RebA_DC were all purified in monomeric form. The interactions between these proteins were examined by far-Western blot assays. The result showed that RebA and RebA_C, but not RebA_DC and RneC, was able to interact with RneN396 (Figure 1D). Since RneN396 does not contain the Zn-link region required for RNase E tetramerization, its interaction with RebA indicates that the tetrameric state of RNase E is not a prerequisite for RebA binding.

Similar to the *E. coli* RNase E, the catalytic region of *Anabaena* RNase E can be further divided into several structurally distinct subregions (i.e., S1, 5’ sensor, RNase H, and DNase I) that are all required for the enzyme activities (Figure 1E). We then examined the interaction between each of these subregions and RebA using bacterial two-hybrid assays. The result showed that interaction occurred only between the 5’ sensor subregion of RNase E and RebA (Figure 1E).

Based on these interaction assays, we concluded that the interaction between RebA and RNase E was mediated by the C domain of RebA and the 5’ sensor domain of RNase E. We also measured the binding affinity between RebA and RneN396 by surface plasmon resonance (SPR) assay (Figure 1F). Their dissociation constant was determined to be 10.3 nM, indicating that the two proteins form a stable complex. The high affinity between RebA and RneN396 allowed us to visualize a clear band of RebA-RneN396 complex in a native gel (Figure 1G). By further analyzing the complex for the molar ratio between them with SDS-PAGE (Figure 1G), we estimated that RneN396 and RebA form a complex with a stoichiometry of 1:1.

### RebA inhibits the binding and the cleavage activities of RNase E

RebA does not contain any domains or motifs that may confer the activities for RNA cleavage or binding, consistent with our results tested with synthetic RNAs in the presence of RebA (Figures 2A, 2B and S2A). However, with 5’ p-LU13-FAM, a 13-nt oligoribonucleotide with a 5’-monophosphate and a 3’ FAM modification that has been used as an RNase E substrate previously (Zhou et al., 2020), we found that RebA when present at more than 2-fold molar excess could strongly interfere with the substrate binding activity of RNase E (Figure 2C), suggesting that it may function as a regulator of the RNase E activity. We thus further tested the RNase E activity in the presence of RebA. RneN412 (1-412 aa) that contains the Zn-link domain for tetramerization was used in most enzymatic assays in this study, as it is easier to purify than the full-length RNase E, while having a similar enzymatic activity (Zhang et al., 2014).

**Figure 2.**
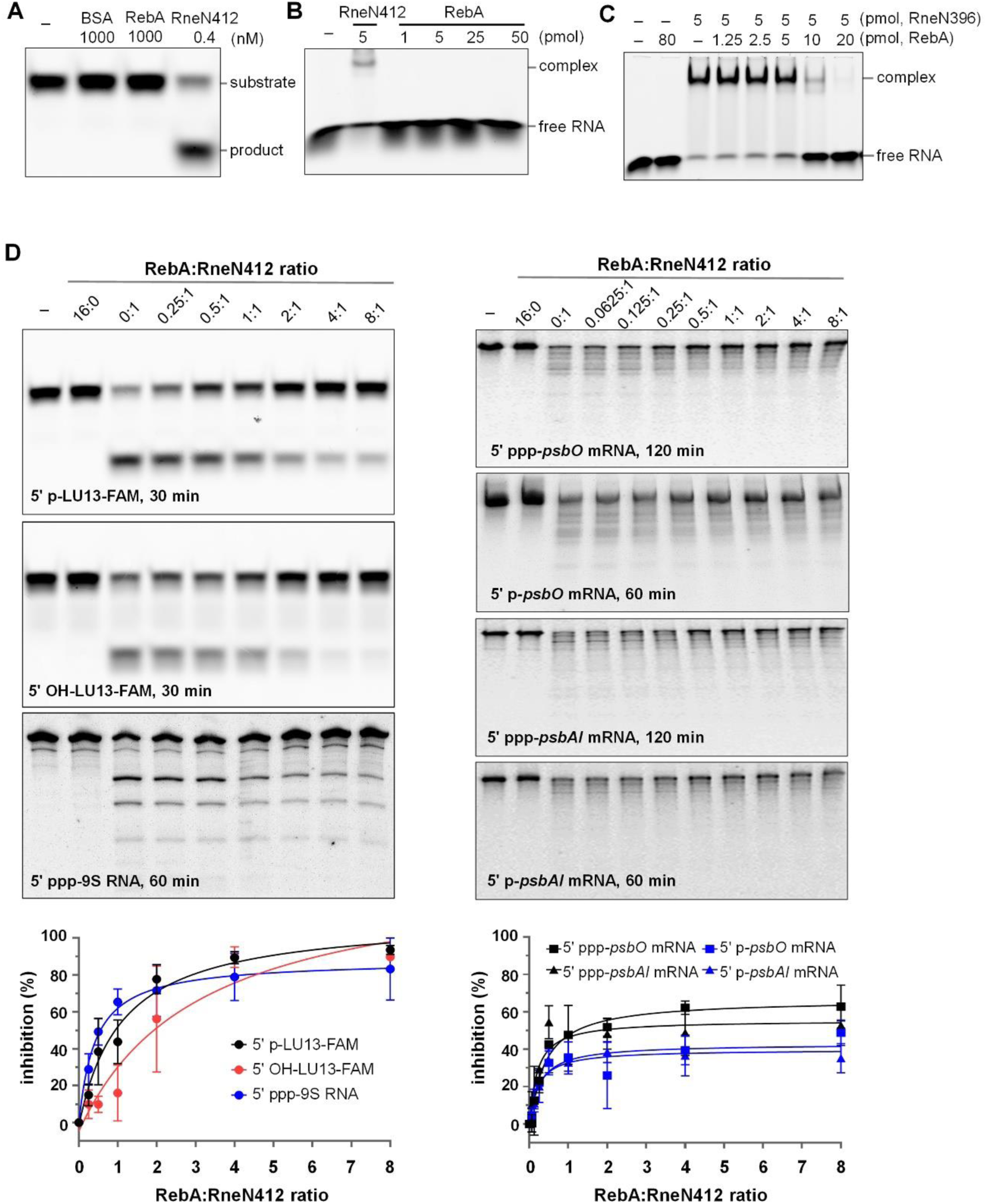
RebA inhibits RNase E activities. **(A)** No cleavage or degradation activities of RebA on the substrate 5’ p-LU13-FAM, a 13-bp RNA (LU13) with a monophosphate at the 5’ end and a 3’ FAM label at the 3’ end. RNA was used at 50 nM. BSA and RneN412 were used as a negative control and a positive control, respectively. The leftmost lane (–) contained RNA only. After 30 minutes of incubation, the reactions were stopped and analyzed by urea-PAGE. RebA also showed no cleavage/degradation activities on other substrates tested (Figure S2). **(B)** No binding activities of RebA using 5’ p-LU13-FAM RNA. The RNA was incubated in the binding buffer with RneN412 (positive control) or with increasing amounts of RebA, respectively. One picomole of RNA was used in the binding mixtures. After 30 minutes of incubation, the reactions were analyzed by native-PAGE. **(C)** Effect of RebA on the binding activity of RneN396 on 5’ p-LU13-FAM. The assay condition was similar to that presented for panel B, except that RebA and RneN396 were incubated on ice for 30 minutes prior to the addition of RNA. **(D)** The cleavage of various RNA substrates by RneN412 in the presence of RebA. 5’ OH-LU13-FAM: similar to 5’ p-LU13-FAM but with a hydroxyl group at the 5’ end; 5’ ppp-9S RNA: *E. coli* 9S RNA with a triphosphorylated 5’ end; *psbAI* mRNA and *psbO* mRNA: mRNAs encoding the photosystem II components D1 and PsbO, respectively. RneN412 were used at 0.4 nM for 5’ p-LU13-FAM, 10 nM for 5’ OH-LU13-FAM, and 200 nM for the other substrates, respectively. All the reactions were performed at 30°C for indicate time periods. For the reactions containing 5’ p-LU13-FAM or 5’ OH-LU13-FAM, 50 nM substrate was included, and 15% urea-PAGE was used to analyze the cleavage products. For the reactions containing 9S RNA, *psbA1* mRNA or *psbO* mRNA, 50 ng/μL substrate and 6% urea-PAGE were used, and gels were stained with GelRed after electrophoresis. The intensities the substrate bands from all three replicative assays were quantified using Image J and used to make the plots.

It is well known that RNase E cleaves many RNA substrates preferentially when they have a 5’-monophosphate end, and such preference involves the recognition of 5’ monophosphate by the 5’ senor domain (Mackie, 1998; Callaghan et al., 2005). We thus tested first with the substrate 5’ p-LU13-FAM. As expected, RneN412 alone efficiently cleaved most of 5’ p-LU13-FAM within 30 minutes (Figure 2D). In the presence of RebA, however, the enzyme activity was significantly inhibited, and this inhibition became stronger with increasing amount of RebA (Figure 2D). According to the inhibition curve, half-inhibition requires a RebA to RneN412 ratio of 1:1. When the ratio reached 4:1, the activity dropped by about 90%.

We further examined the activity of RneN412 in the presence of RebA on two substrates that do not have a 5’-monophosphate for recognition by the 5’ senor domain: 5’ OH-LU13-FAM that is similar to 5’ p-LU13-FAM, but with a 5’-hydroxyl end, and *E. coli* 9S RNA (246 nt) that has a 5’-triphosphate end. These two substrates normally do not have a 5’-monophosphate for recognition by the 5’ senor domain, and are cleaved by RNase E with lower efficiency (Cormack and Mackie, 1992; Kime et al., 2010). Surprisingly, the activity of RneN412 on them was also inhibited by RebA, and the inhibition curves were similar to that for 5’ p-LU13-FAM (Figure 2D).

Furthermore, we tested the inhibition of RebA on RNase E with two *in vitro* transcribed *Anabaena* mRNAs, *psbAI* mRNA (1209 nt) and *psbO* mRNA (1314 nt), which encode the photosystem II components D1 and PsbO, respectively. We chose these two mRNAs because their 5’ boundaries had been clearly defined (Mitschke et al., 2011) and their 3’ boundaries could be predicted with high confidence due to the presence of typical Rho-independent terminators (Figure S2B). These mRNAs, longer than the substrates used above, are expected to have more complex structures. Both mRNAs, either with a 5’-ppp end or a 5’-p end, could be processed by RneN412, and this activity was again inhibited by the presence of RebA, although the maximal inhibition efficiency achieved for these long substrates was lower than those with shorter ones (Figure 2D).

Since the catalytic activity of RNase E is conserved in bacteria, we wondered whether RebA had a cross-species activity. Thus, we tested the cleavage activity of the catalytic domain of *E. coli* RNase E (EcRneN529, 1-529 aa) using 5’ p-LU13-FAM and 9S RNA as substrates in the presence of RebA (Figure S3A). It showed that RebA had no effect on the activity of EcRneN529, consistent with the lack of interaction between RebA and EcRneN529 in bacterial two-hybrid assays (Figure S3B).

Taken together, we conclude that RebA is an inhibitor specifically for cyanobacterial RNase E, and it by interacting with the 5’ sensor domain inhibits the activity of RNase E on a large spectrum of RNA substrates, irrespective of the nature of their 5’ ends.

### RebA plays a role in cell morphology control

To explore the physiological function of RebA, we created a *rebA* deletion mutant (Δ*rebA*) by markerless deletion using a CRISPR/Cpf1-based gene editing system, and an overexpression strain of *rebA* (OE-RebA) by expressing an extra copy of *rebA* controlled by the strong promoter of *rbcL* (P*rbcL*) on a replicative plasmid (Figures S4A to S4D). The growth of Δ*rebA* and OE-RebA in liquid medium and on agar plate were compared with that of the WT. While Δ*rebA* grew only slightly slower than the WT, the overexpression strain OE-RebA grew poorly, especially on agar plate (Figures 3A and 3B). Microscopic images showed that cells of Δ*rebA* appeared significantly shorter than WT cells (Figures 3C and 3D). Note that the extent of morphological changes is dependent on growth conditions: cells were shortened more dramatically in fast-growing cultures; however, cell morphology became closer to that of WT when cells were growing slowly (for example when the culture reached high cell densities or was illuminated under low light intensity) (Figures S4E and S4F). By contrast, cells of OE-RebA were significantly longer and wider than those of WT (Figures 3C and 3D), and this phenotype was not influenced by growth conditions. These results suggested that overexpression of RebA affected negatively cell division or favored lateral peptidoglycan synthesis, leading to elongated cells, while the deletion of *rebA* resulted in an opposite phenotype.

**Figure 3.**
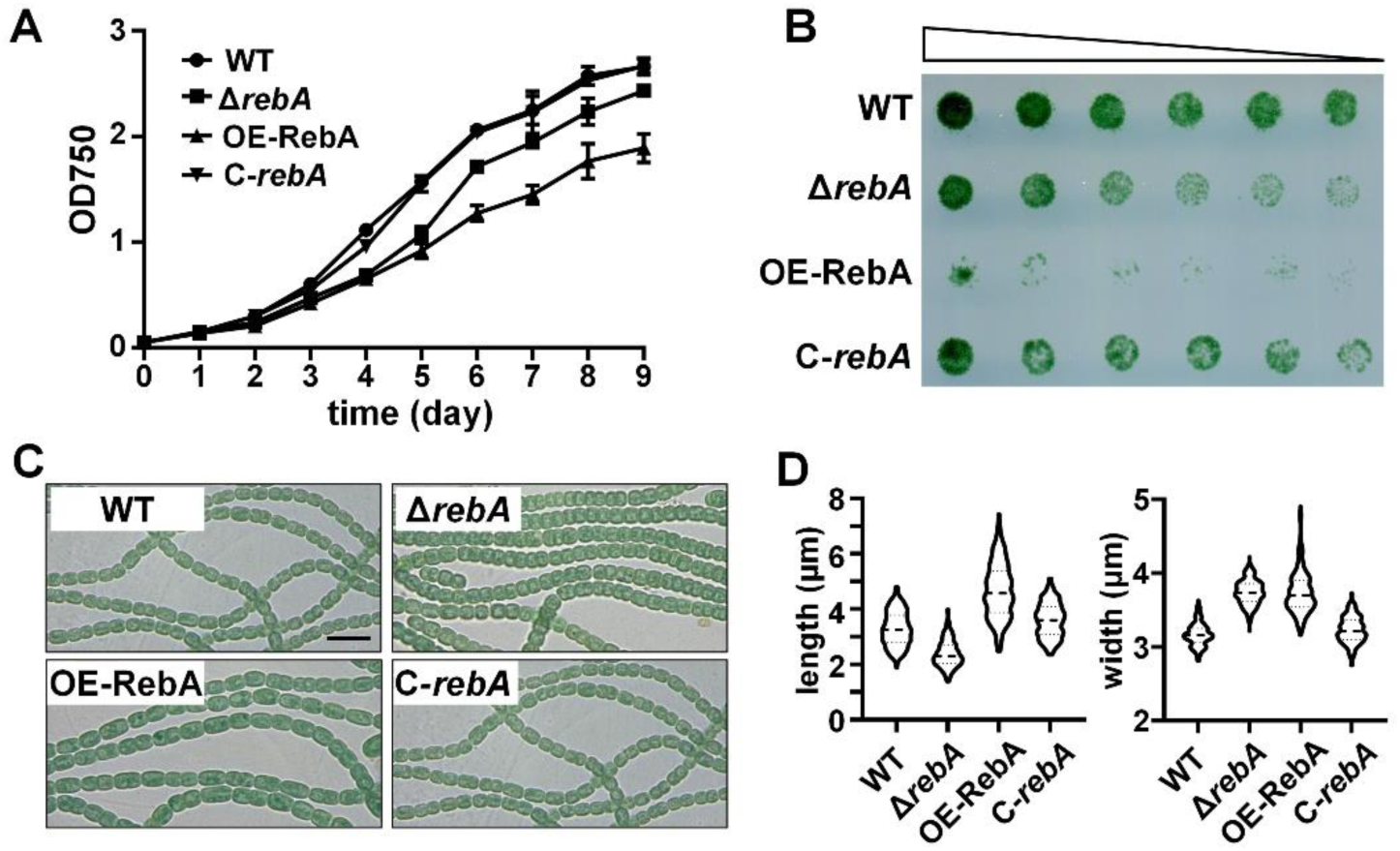
Effect of RebA level on the growth and morphology of *Anabaena* cells. **(A)** Growth of WT (wild type), Δ*rebA*, OE-RebA (RebA overexpression stain using the promoter P*rbcL*), C-*rebA* (Δ*rebA*-complemented strain) in BG11 liquid medium. The optical densities at 750 nm (OD_750_) at different time points are shown as means with standard deviation (n = 3). **(B)** Growth of the same strains as in panel A on solid media. The cultures at OD700 = 0.5 were serially diluted and spotted on the agar plates. The plate was pictured 8 days after incubation. **(C)** Microscopic images of the filaments of different strains. Samples were taken from fresh liquid cultures at OD700 ≈ 0.3. Scale bars:10 μm. **(D)** Distribution of the cell lengths and the cell widths of the samples shown in panel C. A total of 256 cells for each sample was measured.

### RNase E and RebA control cell morphology in an opposite manner

Given that RebA interacted with, and inhibited the activity of RNase E, the phenotypes of Δ*rebA* and OE-RebA could be attributed to altered cellular activities of RNase E. We tested this assumption by modulating the RNase E levels in cells. We initially attempted to create a *rne* deletion strain but failed, implying that the gene could be essential in *Anabaena*, as suggested by similar studies in other cyanobacteria (Horie et al., 2007; Cameron et al., 2015; Cavaiuolo et al., 2020; Hoffmann et al., 2021). We therefore made a conditional mutant of *rne* (CT-*rne*), in which the upstream region (555 bp) of the *rne* ORF was replaced with the CT promoter, inducible by copper and theophylline (Figure 4A). The strain CT-*rne* was maintained in the presence of 0.3 μM Cu^2+^ and 1 mM theophylline. Under such conditions, CT-*rne* showed optimal growth (Figure S5A). When cells of CT-*rne* were transferred to BG11 medium without inducers, the cellular level of RNase E gradually decreased and became nearly undetectable after 5 days (Figure 4B). At the same time point, cells lost viability (Figure S5B), providing experimental evidence for the essential functions of RNase E. By observation under a microscope of CT-*rne* cells, we found that the level of RNase E also had an effect on cell morphology. When RNase E was overexpressed in the presence of high concentrations of inducers (3 μM Cu^2+^ and 2 mM theophylline), cells became shorter and wider, a phenotype similar to that of the Δ*rebA* mutant. In contrast, when RNase E levels decreased following the removal of the inducers, cells became longer, similar to that of the *rebA* overexpression strain OE-RebA (Figures 4C to 4E). Thus, RNase E and RebA both regulate cell morphology, but with opposing functions.

**Figure 4.**
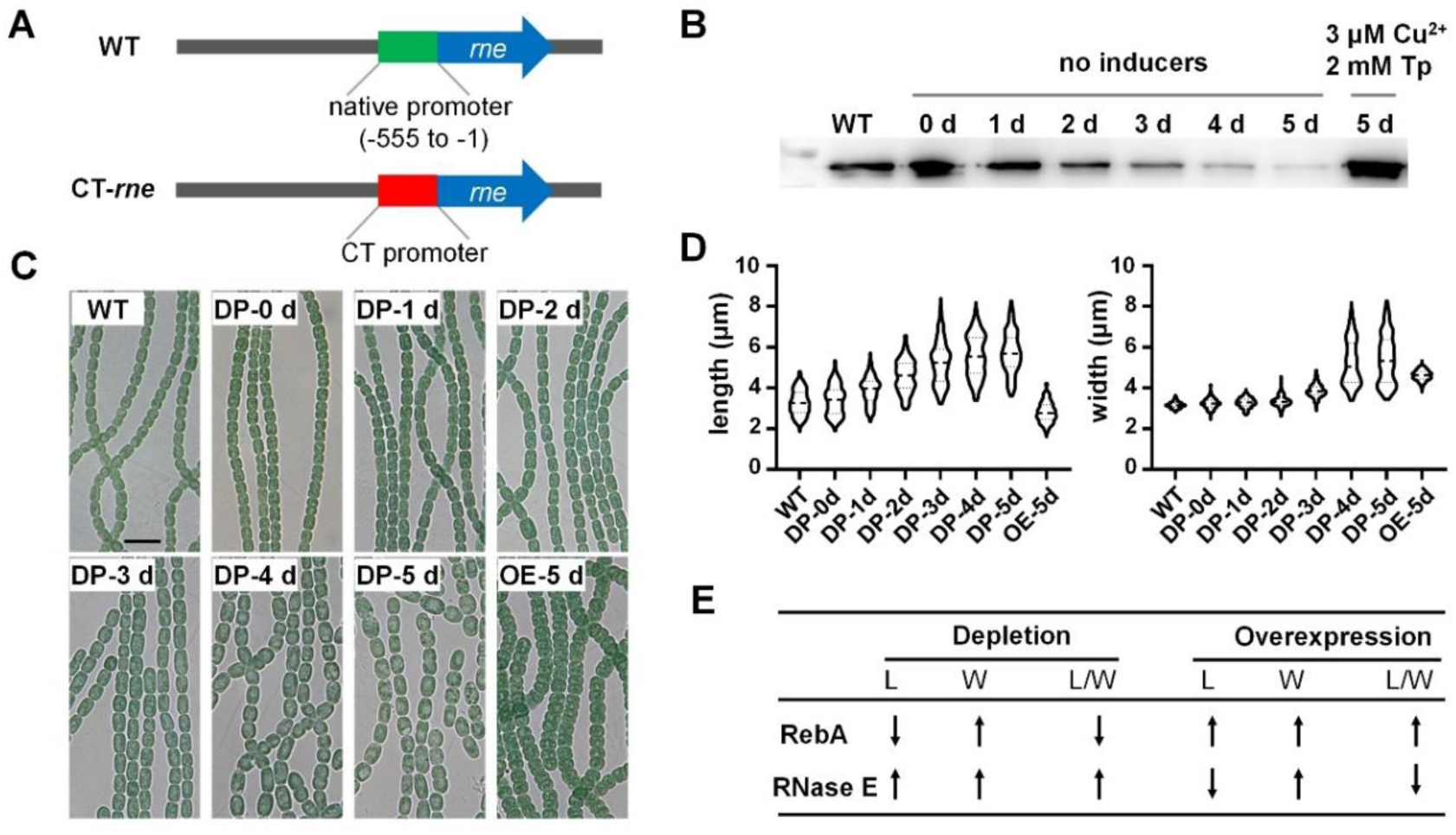
Characterization of a conditional mutant of *rne* (CT-*rne*) and the effects of different levels of RNase E on cell morphology. **(A)** Schematic representation of the genotype of *Anabaena* wild type strain (WT) and that of CT-*rne*. In CT-*rne*, the promoter of *rne* was replaced with the CT promoter, inducible by Cu^2+^ and theophylline (Tp) (Zhou et al., 2020). **(B)** Levels of RNase E in CT-*rne* cells following removal of the inducers or addition of high amounts of inducers in the media. CT-*rne* cells pre-cultured in the presence of 0.3 μM Cu^2+^ and 1 mM theophylline, conditions allowing cells to display a morphology similar to that of WT, were transferred into a medium without Cu^2+^ and Tp for RNase E depletion, or into a medium supplied with 3 μM Cu^2+^ and 2 mM Tp for RNase E overproduction. Cells were collected at the indicated time points for Western blot analysis using antibodies against RNase E. WT cells grown in BG11 was used as a control. Each lane contained 50 μg of cellular proteins. **(C)** Microscopic images of the same samples as those used in panel B. “DP” and “OE” are abbreviations for “depletion” and “overexpression” of RNase E, respectively. Scale bars: 10 μm. **(D)** Distribution of cell lengths and cell widths of the samples shown in panel C. A total of 256 cells from each sample was measured to make the plots. **(E)** Summary of the relationships between cell morphological changes and the depletion or overexpression of RebA and RNase E. L: cell length; W: cell width; L/W: the ratio of cell length to width.

In the genomes of *Anabaena* and many other cyanobacteria, the *rne* gene is immediately upstream of *rnhB* that encodes RNase HII, and the two genes are likely cotranscribed. The phenotypes of CT-*rne* could thus be the result of altered expression of the downstream *rnhB* gene. To rule out this possibility, we constructed a deletion mutant (Δ*rnhB*) and an overexpression strain (OE-RnhB) of *rnhB*. Both strains could grow well in BG11 without apparent morphological changes of the cells (Figure S6), indicating that the observed phenotypes of CT-*rne* were indeed due to the altered expression of *rne*.

### Interaction-dependent regulation of the RNase E activity by RebA is required for maintenance of cell morphology

Since RebA could interact with and inhibit the activity of RNase E, we sought to determine the genetic significance of their interaction. For this purpose, we initially screened for the mutations in RebA that could disrupt RebA-RNase E interaction (Figure S7). In the first-round of mutagenesis, most residues in RebA_C were individually mutated into a charged residue (glutamate or arginine) to maximize the effects of the mutation, followed by bacterial two hybrid assays to check their interaction with RneN396. Those residues whose mutation led to a significantly decreased interaction were subjected to a second-round of mutagenesis in which each residue was replaced by a small, neutral residue (alanine or serine), followed by bacterial two-hybrid assays again. Finally, 11 residues whose mutations would significantly disrupt the interaction in both rounds of mutagenesis (Figure 5A). For some of the residues, we expressed and purified the recombinant RebA mutants with alanine substitution, and examined their inhibitory activities on RNase E. As expected, these RebA derivatives had a greatly reduced capacity of inhibition on the RNase E activity (Figure 5B). Next, the genetic effect of these mutations in RebA was examined following ectopic expression in WT. While overexpression of the WT RebA inhibited growth and led to the formation of elongated cells similar to that caused by RNase E depletion, that of its interaction-defective mutants had much weaker effects on the growth and showed wild-type-like cell morphology (Figures 5C to 5E). These results all together indicate that the inhibition of RNase E by RebA is required for a balanced RNase E activity *in vivo*, and this regulation is dependent on their physical interaction.

**Figure 5.**
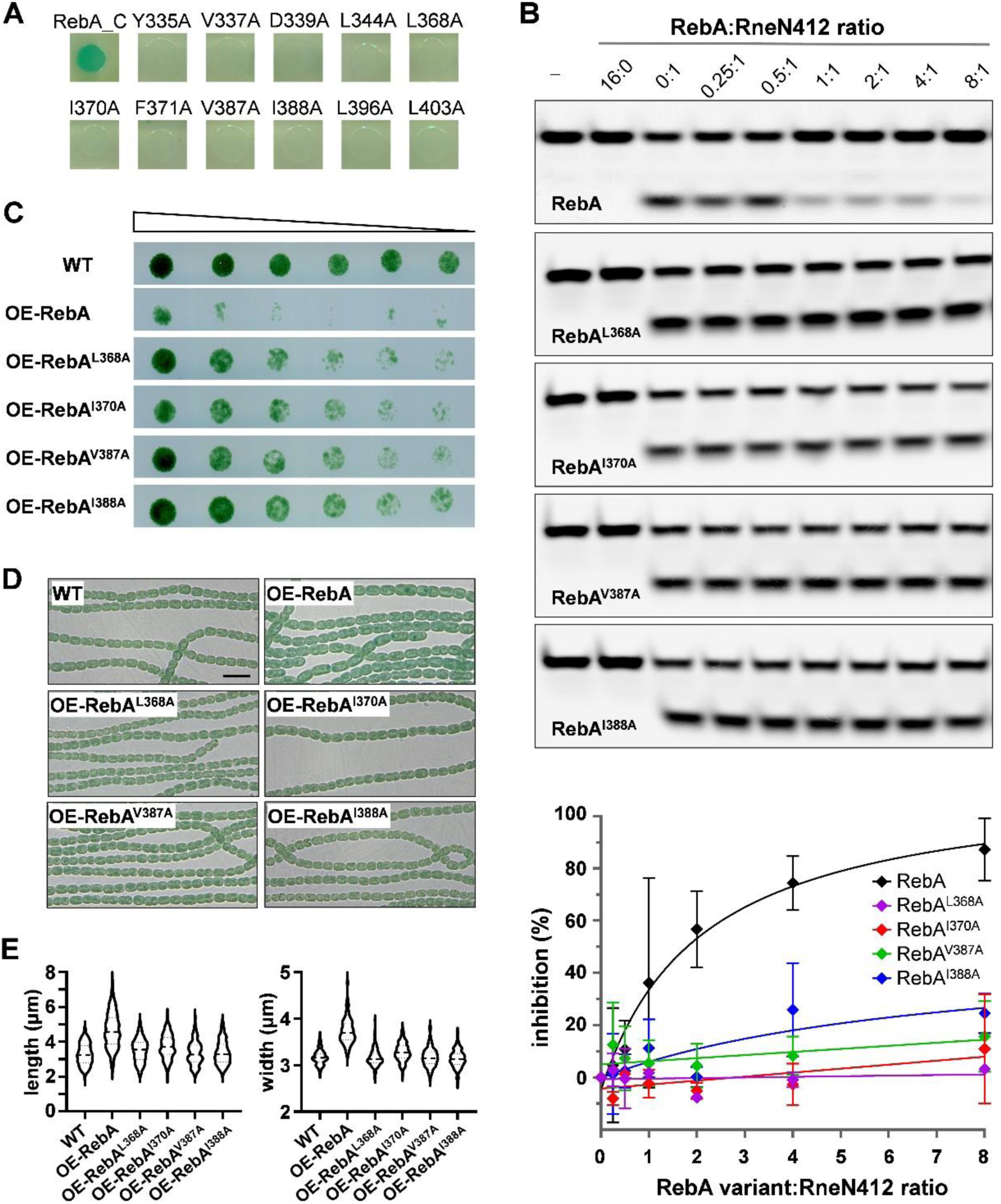
Significance of RebA-RNase E interaction. **(A)** Identification of key residues within RebA_C involved in the interaction with RNase E. Each residue was replaced by alanine, and the interaction was assessed by using bacterial two hybrid assays. Only those mutations that led to significantly decreased interaction were shown here after an initial screening. The result of all mutations within RebA_C was shown in Figure S7. WT RebA_C was used as a positive control. The activity of RNase E on 5’ p-LU13-FAM in the presence of mutant RebA RebA^L368A^, RebA^I370A^, RebA^V387A^ or RebA^I388A^. WT RebA was used as a control. The reaction conditions and data analysis were the same as those for Figure 2D. **(B)** Growth of strains with overproduction of RebA interaction-defective mutants on BG11 agar plates. WT strain and a WT RebA overproduction strain were included as controls. The experiment was performed as described in Figure 3. **(D)** Microscopic images of filaments of different strains grown in BG11 liquid media. Samples were taken from fresh cultures at OD700 ≈ 0.3. Scale bar: 10 μm. **(E)** Distribution of cell lengths and cell widths of the samples shown in the panel D. A total of 256 cells from each sample was measured.

To further confirm the relationship between RebA and RNase E, we examined the effects of *rebA* inactivation while varying the levels of RNase E in the cells. The *rebA* gene was thus deleted from the chromosome in the CT-*rne* strain, leading to strain CT-*rne*::Δ*rebA*. The CT-*rne*::Δ*rebA* strain displayed constantly a shorter-cell phenotype in the presence 0.3 μM copper and 1 mM theophylline (Figure S5C), inducer concentrations that allowed RNase E to be produced at a level without causing cell shape changes in CT-*rne* (Figure 4C). Under such conditions, the absence of RebA in CT-*rne::*Δ*rebA* mimicked therefore RNase E overexpression. Following RNase E depletion by removal of the inducers from the medium, the cells of CT-*rne*::Δ*rebA* changed their shape and maintained viability for significantly longer time than the cells of CT-*rne* (Figure S5C), demonstrating that *rebA* inactivation could slow down the effects of RNase E depletion. These results suggest that a proper level of RNase E activity in the cells is necessary for cell growth and morphology maintenance, since the absence of RebA inhibitor leads to an enhanced activity of RNase E *in vivo*, and partly suppresses the lethal effect caused by decreased levels of RNase E.

## DISCUSSION

RNase E is widely present in bacteria and it plays a central role in RNA metabolism. Due to its broad substrate specificity, its activity must be under check according to changes of internal or external conditions. Yet, only a few regulators of RNase E have been described (Lee *et al*., 2003; Singh et al., 2009; Górna et al., 2010; Moore et al., 2021), and most of them are present only in a limited number of bacteria, suggesting that distinct regulation mechanisms may have been evolved in different organisms. Here, we report that RebA, a protein universally present in cyanobacteria, could inhibit the activity of RNase E. RebA was co-isolated with RNase E from *Anabaena* cell lysate by co-immunoprecipitation. This interaction was further confirmed by bacterial two hybrid assays, and the identification of mutations that disrupted their interaction associated with the expected phenotypes. High affinity interaction occurred between the conserved C-terminal region of RebA and the 5’-sensor domain within the catalytic region of RNase E. RebA showed no RNA-binding or cleavage/degradation activities; it however strongly inhibits the substrate binding and cleavage activities of RNase E. Furthermore, the cellular levels of RebA and RNase E were shown to have opposite effects on cell morphology, demonstrating the inhibitory role of RebA on the RNase E activity *in vivo*. Taken together, we conclude that RebA is an inhibitor of RNase E in *Anabaena*. By varying the relative amounts of RebA and RNase E, we were able to provide evidence on the importance of a balanced activity of RNase E in *Anabaena*.

The activity of RNase E is regulated in multiple ways, including subcellular localization, transcriptional autoregulation, and formation of RNA degradosome (Jain and Belasco, 1995; Prud’homme-Généreux et al., 2004; Murashko et al., 2012; Zhou et al., 2020). Additionally, RNase E activity can also be regulated by transacting components, such as the *E. coli* proteins of RraA, RraB, ribosomal L4 protein, AmiC (Lee et al., 2003; Gao et al., 2006; Singh et al., 2009; Moore et al., 2021). These regulators mostly associate with the noncatalytic region of RNase E, and either enhancing or disrupting their interactions with RNase E did not severely affect cell growth and RNA metabolism of the corresponding bacteria, suggesting their limited regulatory effects *in vivo*. We found here that RebA acts on RNase E in a distinct mechanism: it binds to the catalytic region of RNase E, and inhibits RNase E activity by interfering with substrate binding (Figure 6A). Thus, the effect of RebA on RNase E activity represents a new regulatory mechanism for RNase E functions. Furthermore, RebA could inhibit the cleavage of various substrates by RNase E *in vitro*, and its overexpression led to severe growth defect, similarly as that caused by RNase E depletion, indicating that the control of RNase E by RebA has a global effect on cellular RNA metabolism.

**Figure 6.**
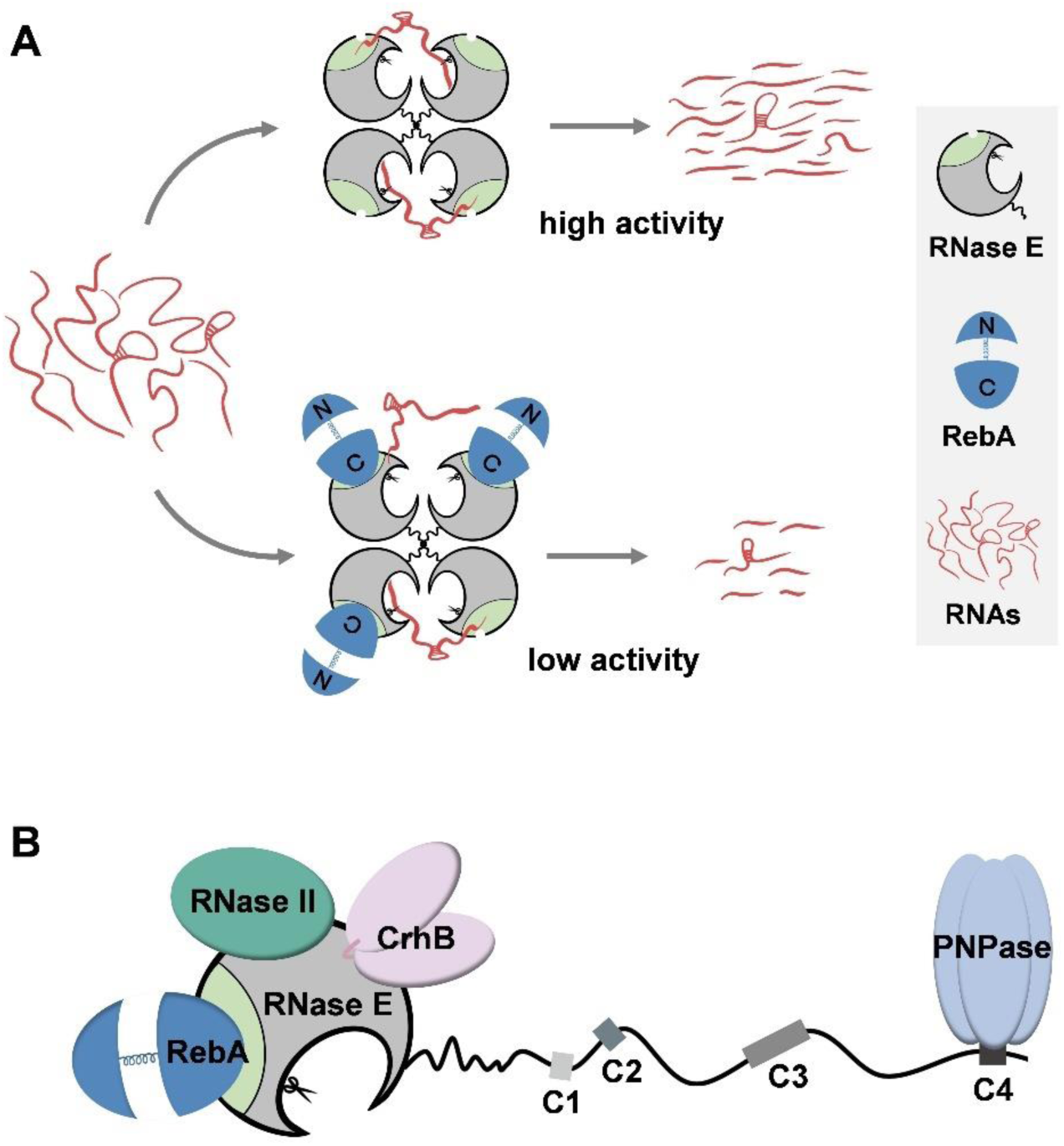
A proposed mechanism of RebA-RNase E interaction, and a proposed RNase E-dependent RNA degradosome in *Anabaena* based on current data. **(A)** Proposed model of RNase E activity regulated by RebA. In the absence of RebA, the subunits in the RNase E tetramer can efficiently bind to RNA substrates, mostly in a 5’-sensor-domain-dependent way. Such a binding ensures a high cleavage activity of RNase E. In contrast, when most or all 5’ sensor domains are bound by RebA, RNase E tetramer binds to substrate less efficiently, resulting in a low cleavage activity. The non-catalytic region of RNase E subunits is not shown in the diagrams for simplification. **(B)** Schematic representation of *Anabaena* degradosome. RebA and PNPase binds to the 5’ sensor domain and the conserved motif C4, respectively. RNase II and CrhB both binds to the catalytic domain of RNase E, but their precise recognition sites on RNase E have not been determined. Only one subunit of the RNase E tetramer is shown for simplicity.

Several substate binding motifs have been identified within both the catalytic and noncatalytic domains of *E. coli* RNase E (Kaberdin et al., 2000; Callaghan et al., 2004; Callaghan et al., 2005; Bandyra et al., 2018). Among them, the 5’-sensor domain within the catalytic region, which has a binding pocket that specifically accommodates the 5’ monophosphate end of RNA, is vital for the binding and the cleavage of 5’-monophosphorylated substrates (Callaghan et al., 2005; Kime et al., 2014). How much this domain contributes to the recognition of other types of substrates has been unclear. We showed here that when the 5’-sensor domain was bound to RebA, the binding and the cleavage of the RNA substrates with 5’-OH, 5’-p or 5’-ppp by RNase E were all inhibited. This result implies that the 5’-sensor domain plays a critical role in the binding and cleavage of all RNA species, regardless of their 5’-end states. The substrate-binding surface within the 5’-sensor domain may be blocked or become inaccessible upon RebA association (Figure 6A).

RebA and RNase E both influence cell morphology of *Anabaena*, and the effect of RebA on cell morphology was mediated by RNase E. Notably, *E. coli* RNase E was shown to have a very similar effect on cell morphology (Goldblum and Apririon, 1981; Tamura et al., 2006; Murashko and Lin-Chao, 2017). In *E. coli*, RNase E can enhance FtsZ translation by degrading the small RNA DicF, which binds to the RBS region of *ftsZ* mRNA and thereby inhibits its translation (Murashko and Lin-Chao, 2017). It remains to be investigated what mechanism occurs in *Anabaena*. Nevertheless, regulation of cell morphology by RNase E in different bacteria suggests a convergent mechanism for the coordination between global RNA metabolism and cell growth.

Based on previous studies, an RNase E-based RNA degradosome, which consists of RNase E, PNPase, RNase II and CrhB, has been suggested to exist in *Anabaena* (Zhang et al., 2022). Here, the low dissociation constant between RebA and RNase E indicates that the two proteins form a stable complex *in vivo*. In this regard, we propose RebA as a new component of the RNA degradosome in *Anabaena* (Figure 6B).

RebA, like RNase E, is encoded by each sequenced cyanobacterial genome. Although RebA homologs from different cyanobacteria have variable lengths; their C-terminal regions, the potential RNase E-interacting region, are highly conserved. Such an observation suggests that the regulation of RNase E by RebA is an ancient and highly conserved mechanism of RNA metabolism control in cyanobacteria. This is supported by a recent finding showing that a RebA homolog and RNase E are present in the same component during the Grid-Seq analysis in *Synechocystis* PCC 6803 (Riediger et al., 2021), a unicellular cyanobacterium phylogenetically distant from *Anabaena* PCC 7120. Our study provides a new control mechanism on gene expression and open the door for our understanding on how RNA metabolism is regulated in different cyanobacterial species in response to changes of physiological or environmental conditions.

## MATERIALS AND METHODS

### Strains and culture conditions

All strains used in this study are listed in Table S1.

The cyanobacterial strains were constructed by transferring the relevant plasmids into the wild type strain of *Anabaena* by conjugation as previously described (Cai and Wolk, 1990; Elhai et al., 1997). Construction of markerless strains using the CRISPR/Cpf1-based genome editing system was according to our previously developed method (Niu et al., 2019). The genotypes of all obtained strains were verified by PCR.

The wild type strain of *Anabaena* and its derivatives were grown in the mineral medium BG11 medium under continuous illumination (30 μmol m^−2^s^−1^) at 30°C with the shaking speed of 180 rpm. When needed, neomycin (50 µg mL^-1^), spectinomycin (5 µg mL^-1^), or streptomycin (2.5 µg mL^-1^) was added into the media. The *rne* conditional mutant strain CT-*rne*, in which the expression of RNase E was induced by the artificial CT promoter (Zhou et al., 2020), was maintained in BG11 supplemented with 0.3 µM CuSO_4_ and 1 mM theophylline.

All *E. coli* strains were grown in LB medium. When needed, kanamycin (50 µg mL^-1^), spectinomycin (100 µg mL^-1^), chloramphenicol (20 µg mL^-1^), or carbenicillin (50 µg mL^-1^) was added into the media.

### Construction of plasmids

All plasmids and related oligonucleotides are listed and additionally described in Tables S2 and S3, respectively.

#### Construction of recombinant protein expression plasmids

The vectors pET28a (Invitrogen), pHStag (GenBank accession number: MK948096; Zhou et al., 2020) and pNStrep (GenBank accession number: OP902607) were used to construct recombinant protein expression plasmids. The plasmid for expressing the full-length RebA protein bearing an N-terminal Strep tag (pNStrep-RebA), which has a codon-optimized ORF of *rebA* cloned in the vector pNStrep, was synthesized by GenScript Biotech Corporation. The plasmids for expressing the RebA variants with single residue mutation were constructed by site-directed mutagenesis using pNStrep-RebA as the parent plasmid. The plasmids expressing the RebA_C and RebA_DC were constructed by amplifying the *rebA* regions from *Anabaena* genomic DNA with the primer pairs Pall1338F901/Pall1338R1269 and Pall1338F1/Pall1338R960 and cloning them into pHSTag via the NdeI-XhoI site, respectively. The plasmid for expressing the catalytic region of *E. coli* RNase E (1-529 aa) was constructed by assembling the insert fragment amplified from the *E. coli* genomic DNA with the primer pair Pb1084F1/Pb1084R1587 and the vector fragment amplified from pET28a with PV_1/PV_2 via seamless cloning. To construct the plasmid for expressing RneN396 (the catalytic region containing the first 396 residues of RNase E), the gene fragment was amplified from *Anabaena* genomic DNA with the primer pair Palr4331F1/Palr4331R1188 was cloned into pET28a via the NdeI and XhoI sites. The plasmids pHTAlr4331, pHTAlr4331N412, pHTAlr4331C, which were used to express the full-length of RNase E, RneN412 (the catalytic region containing the first 412 residues of RNase E) and Rne_C (the non-catalytic region of RNase E), respectively, were described previously (Zhang et al., 2014).

#### Construction of cyanobacterial genome editing plasmids

The CRISPR/Cpf1-based genome editing vectors pCpf1b and pCpf1b-Sp were used to create the plasmids for making genome modified *Anabaena* strains, following the methodology described previously (Niu et al., 2019). The plasmid pICT-*rne* was used to construct a conditional mutant of *rne* in *Anabaena*, in which the native promoter of the *rne* gene was replaced with the artificial and inducible CT promoter. To construct this plasmid, we first made an intermediate plasmid by inserting a fragment of spacer sequence (prepared by annealing the oligonucleotide pair cr_alr4331F26mF/cr_alr4331F26mR) into pCpf1b-sp through the two AarI sites. This plasmid was then linearized with BamHI and BglII. At the same time, two *rne* regions were amplified, respectively, from *Anabaena* genomic DNA with Palr4331F1596m/Palr4331R556m and Palr4331F1b/Palr4331R990, and the CT promoter was amplified from the plasmid pCT (Zhou et al., 2020) using the primer pair PcoquwF/PV_19. The four fragments were assembled into pICT-*rne* via seamless cloning. The plasmids pCpf1b-Δ*rebA* and pCpf1b-Δ*rnhB* were used to create the markerless deletion strains for *rebA* (gene id: all1338) and *rnhB* (gene id: alr4332), respectively. To construct pCpf1b-Δ*rebA*, the BamHI- and BglII-linearized vector pCpf1b-Sp and two *rebA* regions amplified from *Anabaena* genomic DNA with the primer pairs Pall1338F1005m/Pall1338R11 and Pall1338F1247a/Pall1338R2225 were assembled into an intermediate plasmid via seamless cloning. Then this intermediate plasmid was digested with AarI, and ligated with the spacer fragment generated by annealing the oligonucleotide pair cr_all1338R1130F/cr_all1338R1130R, resulting in pCpf1b-Δ*rebA*. The plasmid pCpf1b-Δ*rnhB* was created similarly but with different oligoribonucleotides: *rnhB* regions were amplified with Palr4332F797m/Palr4332R1m and Palr4332F679/Palr4332R1464, and the spacer fragment was prepared with alr4332R379F/cr_alr4332R379R.

#### Construction of replicative plasmids used in *Anabaena*

The plasmid pPrbcL-RebA, which was used to complement the Δ*rebA* mutant and to overexpress RebA protein in the WT, was constructed by assembling the vector fragment amplified from pCT with the primer pair P25TNotI-F/P25TBamHI-R, the promoter region of the *rbcL* gene amplified with PPrbcL-F/PPrbcL-R and the *rebA* ORF amplified from the plasmid pNStrep-RebA with Pall1338bF1c/Pall1338bR1272c via seamless cloning. The plasmids for overexpressing the RebA variants in *Anabaena*, which were based on pPrbcL-RebA, were constructed by site-directed mutagenesis.

#### Construction of plasmids for bacterial two-hybrid assays

The bacterial adenylate cyclase two-hybrid plasmids were constructed using the vectors pKT25a (GenBank accession number: OQ032554) and pUT18Ca (GenBank accession number: OP902608), which were modified from pKT25 and pUT18C (Battesti and Bouveret, 2012), respectively. pKT25a and pUT18Ca have identical cloning site, thus allowing the same fragment to be cloned into both vectors. All the two hybrid plasmids were constructed by seamlessly cloning as previously described (Zhou et al., 2020).

All the constructed plasmids were verified by DNA sequencing.

### Bacterial two-hybrid assays

The two-hybrid system of Bacterial Adenylate Cyclase Two-Hybrid (BATCH) was used to test for protein-protein interactions (Battesti and Bouveret, 2012). Pairs of two-hybrid plasmids were co-transformed into the *cya*^−^ strain BTH101. The transformants were grown on LB plates containing 50 µg/mL ampicillin, 25 µg/mL kanamycin, 0.5 mM IPTG (isopropyl β-D-1-thiogalactopyranoside), and 40 µg/mL X-gal (5-bromo-4-chloro-3-indolyl-ꞵ-D galactopyranoside) in the dark at 30°C for 1–3 days for interaction screening. Transformants that contain plasmids encoding interacting proteins are expected to have colonies with blue color. The strain co-transformed with pKT25-zip and pUT18C-zip was used as a positive control (Battesti and Bouveret, 2012), and that co-transformed with the empty vectors of pKT25a and pUT18Ca was used as a negative control.

### Purification of recombinant proteins

*E. coli* BL21(DE3) was used as the host strain for recombinant protein expression. Proteins with a His tag was purified with the Ni-NTA resin (Genscript) according to the product manual. Proteins with a Strep tag were purified using the Strep-Tactin XT resin (IBA-Lifesciences) according to the product manual. When necessary, the proteins eluted from Ni-NTA resin or Strep-Tactin XT resin were further passed through a size-exclusion column to improve the purity. Purified proteins were finally dialyzed into the storage buffer (50 mM Tris-HCl, 500 mM NaCl, 2 mM EDTA and 25% glycerol, pH 8.0), and stored at −80 °C.

### Synthesis of RNA substrates

The short labelled RNAs 5’ p-LU13-FAM (5’ p-GAGACAGUAUUUG-FAM) and 5’ OH-LU13-FAM (5’ OH-GAGACAGUAUUUG-FAM) were chemically synthesized by Genscript Biotech Corporation. The 5’ triphosphorylated RNAs of 9S RNA, the *psbO* mRNA and *psbAI* mRNA were produced by *in vitro* transcription using the T7 High Yield RNA Transcription Kit (Vazyme). The 5’ monophosphorylated RNAs were prepared by providing five-fold excess of GMP over GTP in the *in vitro* transcription reactions. The DNA templates for the transcription of 9S RNA was prepared as previously described (Zhang et al., 2014). The templates for *psbO* mRNA and *psbA* mRNA transcription were amplified from the genomic DNA of *Anabaena* using the primer pairs of Pall3854F219ma/Pall3854R1095 and Palr4866F65maa/Palr4866R1144a. The transcribed RNAs were purified using RNA purification columns.

### RNase E cleavage activity assays

The cleavage activity of RNase E was measured in the reaction buffer containing 20 mM TrisHCl (pH 8.0), 100 mM NaCl, 5 mM MgCl2, 0.1 mM DTT, 5% glycerol, 0.1% Triton X-100, 1 mg/ml BSA and 2 U/μL Murine RNase inhibitor. To investigate the effect of RebA on RNase E activity, RneN412 was first incubated with indicated amount of RebA in the reaction buffer for 30 min on ice. Subsequently, RNA substrate was added into the mixture at 30°C for indicated time periods. The reaction was stopped by adding an equal volume of 2X RNA loading buffer (95% formamide, 5 mM EDTA, 0.025% SDS, 0.025% bromophenol blue, 0.025% xylene-cyanol). After treatment at 95°C for 5 min, cleavage products in the reaction mixture were separated on 5% (for mRNA substrates) or 15% (for oligoribonucleotide substrates) polyacrylamide gels that contained 7 M urea. For FAM-labeled RNA substrates, the gels were detected with GE Typhoon Trio Imager, while for non-labeled substrates, the gels were stained with GelRed (Biotium) and imaged with a common gel imaging system. The intensity of RNA bands was quantified using Image J.

The inhibitory effect of RebA on RNase E activity was evaluated with the RNAs of 5’ p-LU13-FAM, 5’ OH-LU13-FAM, 9S RNA, *psbO* mRNA and *psbAI* mRNA. The cleavage products were separated on PAGE gels, and the intensities of the gel bands were used to calculate the inhibition efficiency (I) of RebA on the RNase activity using the formula:

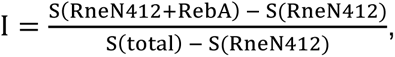

where S(total) is the intensity of the band in the blank control reaction that contains RNA only, S(RneN412+RebA) is the intensity of the uncut substate band when RebA and RneN412 are both added in the reaction, and S(RneN412) is the intensity of the uncut substate band when only RneN412 is included in the reaction.

### RNA-binding assays

RNA-binding assays were carried out in the buffer containing 20 mM TrisHCl (pH 8.0), 200 mM NaCl, 0.1 mM DTT, 2 mM EDTA, 5% glycerol and 1 mg/ml BSA). To test the effect of RebA on the binding activity of RNase E, RneN396 and RebA was first incubated with indicated amounts of RebA in the binding buffer for 30 min on ice, then RNA substrate was added into the mixture for another 30 min of incubation at 30°C. The incubation mixtures were then analyzed on 5% native polyacrylamide gels at 4°C.

### Co-immunoprecipitation

To prepare the magnetic beads covalently coated by RNase E polyclonal antibodies (Anti-Rne), the purified polyclonal antibodies against RNase E was bound to Protein A/G MagBeads (Genscript), followed by cross-linking using dimethyl pimelidate dihydrochloride (Sigma-Aldrich) according to the manufacture’s instruction. For co-immunoprecipitation of cellular proteins associated with RNase E, about 200 ml fresh *Anabaena* cells (OD700 ≈ 0.5) grown in BG11 was collected by centrifugation at 5,000 g for 10 min at room temperature. After washing once with 5 ml buffer W (50 TrisHCl, 150 mM NaCl, pH8.0), cells were resuspended in 1 ml lysis buffer (50 mM TrisHCl, pH8.0, 150 mM NaCl, 0.1% Triton X-100) supplemented with cOmplete™ Protease Inhibitor Cocktail (Roche) and 50 μg/ml of RNase A. Subsequently, the cells were lyzed by FastPrep-24 (MP Biomedicals). Following centrifuge at 12,000 g for 20 minutes, the supernatant of cell lysate was transferred into a 2 ml tube containing 0.2 ml of Anti-Rne beads. After 5 hours of incubation at 4°C with gentle rotation, the beads were collected and washed twice with the lysis buffer. Proteins captured by the Anti-Rne beads were then eluted in 200 μl of 1X SDS-PAGE loading buffer (50 mM TrisHCl, pH 6.8, 50 mM DTT, 2% SDS, 1 mM EDTA, 0.01% bromophenol blue, 5% glycerol). The elute was analyzed by 10% SDS-PAGE, and the protein bands were excised from the gel and identified by mass spectrometry.

### SPR assays

The binding constants between RneN396 and RebA were determined by surface plasmon resonance (SPR) at 25 °C in a Biacore instrument (Cytiva). RneN396 at 2 μM was immobilized covalently on a streptavidin-coated sensor chip in the running buffer of PBS (8 mM Na2HPO4, 1.46 mM KH2PO4, 137 mM NaCl, 2.7 mM KCl, pH7.4). RebA at the concentration series of 0, 31.25, 62.5, 125, 250, 500 and 1,000 nM were injected at the constant flow rate of 500 μL min^-1^. The association was monitored over 180 s and dissociation was monitored over 300 s. The equilibrium dissociation constant was calculated and evaluated using a steady-state affinity program of BIA evaluation software associated with the instrument.

### Western blotting

Conventional Western blotting was performed as previously described (Zhang et al., 2014). The polyclonal antibodies for detecting RNase E and RebA were produced by immunizing rabbits with the purified proteins (FrdBio). The production of the antibodies against FtsZ was described previously (Wang et al., 2021).

Far-Western blotting assay was performed as previously described (Zhang et al., 2014; Zhou et al., 2020). Briefly, to test the interaction between RebA and RNase E regions, RneN396 (1 μg), RneC (1 μg), and RebA (10 ng, used as positive control) were separated in duplicate on two 10% SDS-PAGE gels and electrotransferred onto two PVDF membranes. Both membranes were blocked in PBS-T (PBS supplemented with 0.1% Tween 20) containing 5% skimmed milk for 1 h. Then, one membrane was incubated with 10 μg of RebA in PBS-T containing 1% skimmed milk, while the other was incubated in the same solution but without RebA. Subsequently, both membranes were immunodetected with the antibodies against RebA. The interactions between subregions of RebA (i.e., RebA_C and RebA_DC) and RNase E regions were detected in the same way.

### Quantification and statistical analysis

Statistical analyses were performed using GraphPad Prism. Statistical details for individual experiments, such as number of samples and experimental replicates, have been described in the figure legends and method details.

## ACKNOWLEDGEMENTS

This study was supported by the National Natural Science Foundation of China (Grant No.: 32070037), the Featured Institute Service Project from the Institute of Hydrobiology, the Chinese Academy of Sciences (Grant No.: Y85Z061601), and the Knowledge Innovation Program of Wuhan - Basic Research (Grant No.: 2022020801010142). We thank Tian-Cai Niu for the help in protein purification, and Dr Shaoran Zhang at Huazhong Agricultural University for the help in SPR assay.

## AUTHOR CONTRIBUTIONS

S.-J.L., Y.-Q.Y., G.-M.L. and J.-Y.Z. contributed to experimental works and data analysis. J.-Y.Z. and C.-C.Z. contributed to conceptualization, manuscript writing and editing. J.-Y.Z., C.-C.Z. and W.C. contributed to discussions, resources, and funding acquisition.

## DECLARATION OF INTERESTS

The authors declare no competing interests.

## Notes

### Competing Interest Statement

The authors have declared no competing interest.

## REFERENCES

Aït-Bara S., and Carpousis A.J. (2015) RNA degradosomes in bacteria and chloroplasts: classification, distribution and evolution of RNase E homologs. Mol Microbiol 97: 1021–1135.

Andersson A.F., Lundgren M., Eriksson S., Rosenlund M., Bernander R., and Nilsson P. (2006) Global analysis of mRNA stability in the archaeon *Sulfolobus*. Genome Biol 7: R99.

Babitzke P., and Kushner S.R. (1991) The Ams (altered mRNA stability) protein and ribonuclease E are encoded by the same structural gene of *Escherichia coli*. Proc Natl Acad Sci U S A 88: 1–5.

Bandyra K.J., Wandzik J.M., and Luisi B.F. (2018) Substrate recognition and autoinhibition in the central ribonuclease RNase E. Mol Cell 72: 275–285.e4.

Battesti A., and Bouveret E. (2012) The bacterial two-hybrid system based on adenylate cyclase reconstitution in *Escherichia coli*. Methods 58: 325–334.

Behler J., Sharma K., Reimann V., Wilde A., Urlaub H., and Hess W.R. (2018) The host-encoded RNase E endonuclease as the crRNA maturation enzyme in a CRISPR-Cas subtype III-Bv system. Nat Microbiol 3: 367–377.

Bernstein J.A., Khodursky A.B., Lin P.H., Lin-Chao S., and Cohen S.N. (2002) Global analysis of mRNA decay and abundance in *Escherichia coli* at single-gene resolution using two-color fluorescent DNA microarrays. Proc Natl Acad Sci U S A 99: 9697–9702.

Cai Y.P., and Wolk C.P. (1990) Use of a conditionally lethal gene in *Anabaena* sp. strain PCC 7120 to select for double recombinants and to entrap insertion sequences. J Bacteriol 172: 3138–3145.

Callaghan A.J., Aurikko J.P., Ilag L.L., Günter Grossmann J., Chandran V., Kühnel K., Poljak L., Carpousis A.J., Robinson C.V., Symmons M.F., et al. (2004) Studies of the RNA degradosome-organizing domain of the *Escherichia coli* ribonuclease RNase E. J Mol Biol 340: 965–979.

Callaghan A.J., Marcaida M.J., Stead J.A., McDowall K.J., Scott W.G., and Luisi B.F. (2005) Structure of *Escherichia coli* RNase E catalytic domain and implications for RNA turnover. Nature 437: 1187–1191.

Cameron J.C., Gordon G.C., and Pfleger B.F. (2015) Genetic and genomic analysis of RNases in model cyanobacteria. Photosynth Res 126: 171–183.

Cavaiuolo M., Chagneau C., Laalami S., and Putzer H. (2020) Impact of RNase E and RNase J on global mRNA metabolism in the cyanobacterium *Synechocystis* PCC6803. Front Microbiol 11: 1055.

Chao Y., Li L., Girodat D., Förstner K.U., Said N., Corcoran C., Śmiga M., Papenfort K., Reinhardt R., Wieden H.J., et al. (2017) in Vivo cleavage map illuminates the central role of RNase E in coding and non-coding RNA pathways. Mol Cell 65: 39–51.

Clarke J.E., Kime L., Romero A D., and McDowall K.J. (2014) Direct entry by RNase E is a major pathway for the degradation and processing of RNA in *Escherichia coli*. Nucleic Acids Res 42: 11733–11751.

Cormack R.S., and Mackie G.A. (1992) Structural requirements for the processing of *Escherichia coli* 5 S ribosomal RNA by RNase E in vitro. J Mol Biol 228: 1078–1090.

Elhai J., Vepritskiy A., Muro-Pastor A.M., Flores E., and Wolk C.P. (1997) Reduction of conjugal transfer efficiency by three restriction activities of *Anabaena* sp. strain PCC 7120. J Bacteriol 179: 1998–2005.

Gabr A., Grossman A.R., and Bhattacharya D. (2020) Paulinella, a model for understanding plastid primary endosymbiosis. J Phycol 56: 837–843.

Gao J., Lee K., Zhao M., Qiu J., Zhan X., Saxena A., Moore C.J., Cohen S.N., and Georgiou G. (2006) Differential modulation of *E. coli* mRNA abundance by inhibitory proteins that alter the composition of the degradosome. Mol Microbiol 61: 394–406.

Garrey S.M., Blech M., Riffell J.L., Hankins J.S., Stickney L.M., Diver M., Hsu Y.H., Kunanithy V., and Mackie G.A. (2009) Substrate binding and active site residues in RNases E and G: role of the 5’-sensor. J Biol Chem 284: 31843–31850.

Goldblum K., and Apririon D. (1981) Inactivation of the ribonucleic acid-processing enzyme ribonuclease E blocks cell division. J Bacteriol 146: 128–132.

Górna M.W., Pietras Z., Tsai Y.C., Callaghan A.J., Hernández H., Robinson C.V., and Luisi B.F. (2010) The regulatory protein RraA modulates RNA-binding and helicase activities of the *E. coli* RNA degradosome. RNA 16: 553–562.

Hambraeus G., von Wachenfeldt C., and Hederstedt L. (2003) Genome-wide survey of mRNA half-lives in *Bacillus subtilis* identifies extremely stable mRNAs. Mol Genet Genomics 269: 706–714.

Hoffmann U.A., Heyl F., Rogh S.N., Wallner T., Backofen R., Hess W.R., Steglich C., and Wilde A. (2021) Transcriptome-wide in vivo mapping of cleavage sites for the compact cyanobacterial ribonuclease E reveals insights into its function and substrate recognition. Nucleic Acids Res 49: 13075–13091.

Horie Y., Ito Y., Ono M., Moriwaki N., Kato H., Hamakubo Y., Amano T., Wachi M., Shirai M., and Asayama M. (2007) Dark-induced mRNA instability involves RNase E/G-type endoribonuclease cleavage at the AU-box and *SD* sequences in cyanobacteria. Mol Genet Genomics 278: 331–346.

Huang H., Liao J., and Cohen S.N. (1998) Poly(A)-and poly(U)-specific RNA 3’ tail shortening by *E. coli* ribonuclease E. Nature 391: 99–102.

Jain C., and Belasco J.G. (1995) RNase E autoregulates its synthesis by controlling the degradation rate of its own mRNA in *Escherichia coli*: unusual sensitivity of the rne transcript to RNase E activity. Genes Dev 9: 84–96.

Kaberdin V.R., Miczak A., Jakobsen J.S., Lin-Chao S., McDowall K.J., and von Gabain A. (1998) The endoribonucleolytic N-terminal half of *Escherichia coli* RNase E is evolutionarily conserved in *Synechocystis* sp. and other bacteria but not the C-terminal half, which is sufficient for degradosome assembly. Proc Natl Acad Sci U S A 95: 11637–11642.

Kaberdin V.R., Walsh A.P., Jakobsen T., McDowall K.J., and von Gabain A. (2000) Enhanced cleavage of RNA mediated by an interaction between substrates and the arginine-rich domain of *E. coli* ribonuclease E. J Mol Biol 301: 257–264.

Kime L., Clarke J.E., Romero A D., Grasby J.A., and McDowall K.J. (2014) Adjacent single-stranded regions mediate processing of tRNA precursors by RNase E direct entry. Nucleic Acids Res 42: 4577–4589.

Kime L., Jourdan S.S., Stead J.A., Hidalgo-Sastre A., and McDowall K.J. (2010) Rapid cleavage of RNA by RNase E in the absence of 5’ monophosphate stimulation. Mol Microbiol 76: 590–604.

Lee K., Zhan X., Gao J., Qiu J., Feng Y., Meganathan R., Cohen S.N., and Georgiou G. (2003) RraA. a protein inhibitor of RNase *E* activity that globally modulates RNA abundance in *E. coli*. Cell 114: 623–634.

Mackie G.A. (1998) Ribonuclease E is a 5’-end-dependent endonuclease. Nature 395: 720–723.

Mackie G.A. (2013) RNase E: at the interface of bacterial RNA processing and decay. Nat Rev Microbiol 11: 45–57.

McDowall K.J., Kaberdin V.R., Wu S.W., Cohen S.N., and Lin-Chao S. (1995) Site-specific RNase E cleavage of oligonucleotides and inhibition by stem-loops. Nature 374: 287–290.

McDowall K.J., Lin-Chao S., and Cohen S.N. (1994) A+U content rather than a particular nucleotide order determines the specificity of RNase E cleavage. J Biol Chem 269: 10790–10796.

Miczak A., Kaberdin V.R., Wei C.L., and Lin-Chao S. (1996) Proteins associated with RNase E in a multicomponent ribonucleolytic complex. Proc Natl Acad Sci U S A 93: 3865–3869.

Mitschke J., Vioque A., Haas F., Hess W.R., and Muro-Pastor A.M. (2011) Dynamics of transcriptional start site selection during nitrogen stress-induced cell differentiation in *Anabaena* sp. PCC7120. Proc Natl Acad Sci U S A 108: 20130–20135.

Moore C.J., Go H., Shin E., Ha H.J., Song S., Ha N.C., Kim Y.H., Cohen S.N., and Lee K. (2021) Substrate-dependent effects of quaternary structure on RNase E activity. Genes Dev 35: 286–299.

Murashko O.N., and Lin-Chao S. (2017) *Escherichia coli* responds to environmental changes using enolasic degradosomes and stabilized DicF sRNA to alter cellular morphology. Proc Natl Acad Sci U S A 114: E8025–E8034.

Murashko O.N., Kaberdin V.R., and Lin-Chao S. (2012) Membrane binding of *Escherichia coli* RNase E catalytic domain stabilizes protein structure and increases RNA substrate affinity. Proc Natl Acad Sci U S A 109: 7019–7024.

Niu T.C., Lin G.M., Xie L.R., Wang Z.Q., Xing W.Y., Zhang J.Y., and Zhang C.C. (2019) Expanding the potential of CRISPR-Cpf1-Based genome editing technology in the cyanobacterium *Anabaena* PCC 7120. ACS Synth Biol 8: 170–180.

Ono M., and Kuwano M. (1979) A conditional lethal mutation in an *Escherichia coli* strain with a longer chemical lifetime of messenger RNA. J Mol Biol 129: 343–357.

Prud’homme-Généreux A., Beran R.K., Iost I., Ramey C.S., Mackie G.A., and Simons R.W. (2004) Physical and functional interactions among RNase E, polynucleotide phosphorylase and the cold-shock protein, CsdA: evidence for a ‘cold shock degradosome’. Mol Microbiol 54: 1409–1421.

Py B., Higgins C.F., Krisch H.M., and Carpousis A.J. (1996) A DEAD-box RNA helicase in the *Escherichia coli* RNA degradosome. Nature 381: 169–172.

Richards J., and Belasco J.G. (2019) Obstacles to scanning by RNase E govern bacterial mRNA lifetimes by hindering access to distal cleavage sites. Mol Cell 74: 284–295.e5.

Riediger M., Spät P., Bilger R., Voigt K., Maček B., and Hess W.R. (2021) Analysis of a photosynthetic cyanobacterium rich in internal membrane systems via gradient profiling by sequencing (Grad-seq). Plant Cell 33: 248–269.

Sakurai I., Stazic D., Eisenhut M., Vuorio E., Steglich C., Hess W.R., and Aro E.M. (2012) Positive regulation of *psbA* gene expression by cis-encoded antisense RNAs in *Synechocystis* sp. PCC 6803. Plant Physiol 160: 1000–1010.

Shih P.M., Wu D., Latifi A., Axen S.D., Fewer D.P., Talla E., Calteau A., Cai F., Tandeau de Marsac N., Rippka R., et al. (2013) Improving the coverage of the cyanobacterial phylum using diversity-driven genome sequencing. Proc Natl Acad Sci U S A 110: 1053–1058.

Singh D., Chang S.J., Lin P.H., Averina O.V., Kaberdin V.R., and Lin-Chao S. (2009) Regulation of ribonuclease E activity by the L4 ribosomal protein of *Escherichia coli*. Proc Natl Acad Sci U S A 106: 864–869.

Stanier R.Y., and Cohen-Bazire G. (1977) Phototrophic prokaryotes: the cyanobacteria. Annu Rev Microbiol 31: 225–274.

Stead M.B., Marshburn S., Mohanty B.K., Mitra J., Pena Castillo L., Ray D., van Bakel H., Hughes T.R., and Kushner S.R. (2011) Analysis of *Escherichia coli* RNase E and RNase III activity in vivo using tiling microarrays. Nucleic Acids Res 39: 3188–3203.

Steglich C., Lindell D., Futschik M., Rector T., Steen R., and Chisholm S.W. (2010) Short RNA half-lives in the slow-growing marine cyanobacterium *Prochlorococcus*. Genome Biol 11: R54.

Tamura M., Lee K., Miller C.A., Moore C.J., Shirako Y., Kobayashi M., and Cohen S.N. (2006) RNase E maintenance of proper FtsZ/FtsA ratio required for nonfilamentous growth of *Escherichia coli* cells but not for colony-forming ability. J Bacteriol 188: 5145–5152.

Van den Bossche A., Hardwick S.W., Ceyssens P.J., Hendrix H., Voet M., Dendooven T., Bandyra K.J., De Maeyer M., Aertsen A., Noben J.P., et al. (2016) Structural elucidation of a novel mechanism for the bacteriophage-based inhibition of the RNA degradosome. Elife 5: e16413.

Wang L., Niu T.C., Valladares A., Lin G.M., Zhang J.Y., Herrero A., Chen W.L., and Zhang C.C. (2021) The developmental regulator PatD modulates assembly of the cell-division protein FtsZ in the cyanobacterium *Anabaena* sp. PCC 7120. Environ Microbiol 23: 4823–4837.

Yan H., Qin X., Wang L., and Chen W. (2020) Both enolase and the DEAD-Box RNA helicase CrhB can form complexes with RNase E in *Anabaena* sp. strain PCC 7120. Appl Environ Microbiol 86: e00425–20.

Zhang J.Y., Deng X.M., Li F.P., Wang L., Huang Q.Y., Zhang C.C., and Chen W.L. (2014) RNase E forms a complex with polynucleotide phosphorylase in cyanobacteria via a cyanobacterial-specific nonapeptide in the noncatalytic region. RNA 20: 568–579.

Zhang J.Y., Hess H.R., and Zhang C.C. (2022) “Life is short, and art is long”: RNA degradation in cyanobacteria and model bacteria. mLife 1: 21–39.

Zhou C., Zhang J., Hu X., Li C., Wang L., Huang Q., and Chen W. (2020) RNase II binds to RNase E and modulates its endoribonucleolytic activity in the cyanobacterium *Anabaena* PCC 7120. Nucleic Acids Res 48: 3922–3934.

