## Supplementary Figures S1-S7 and Tables S1-S3 for "A conserved protein inhibitor brings under check the activity of RNase E in cyanobacteria"

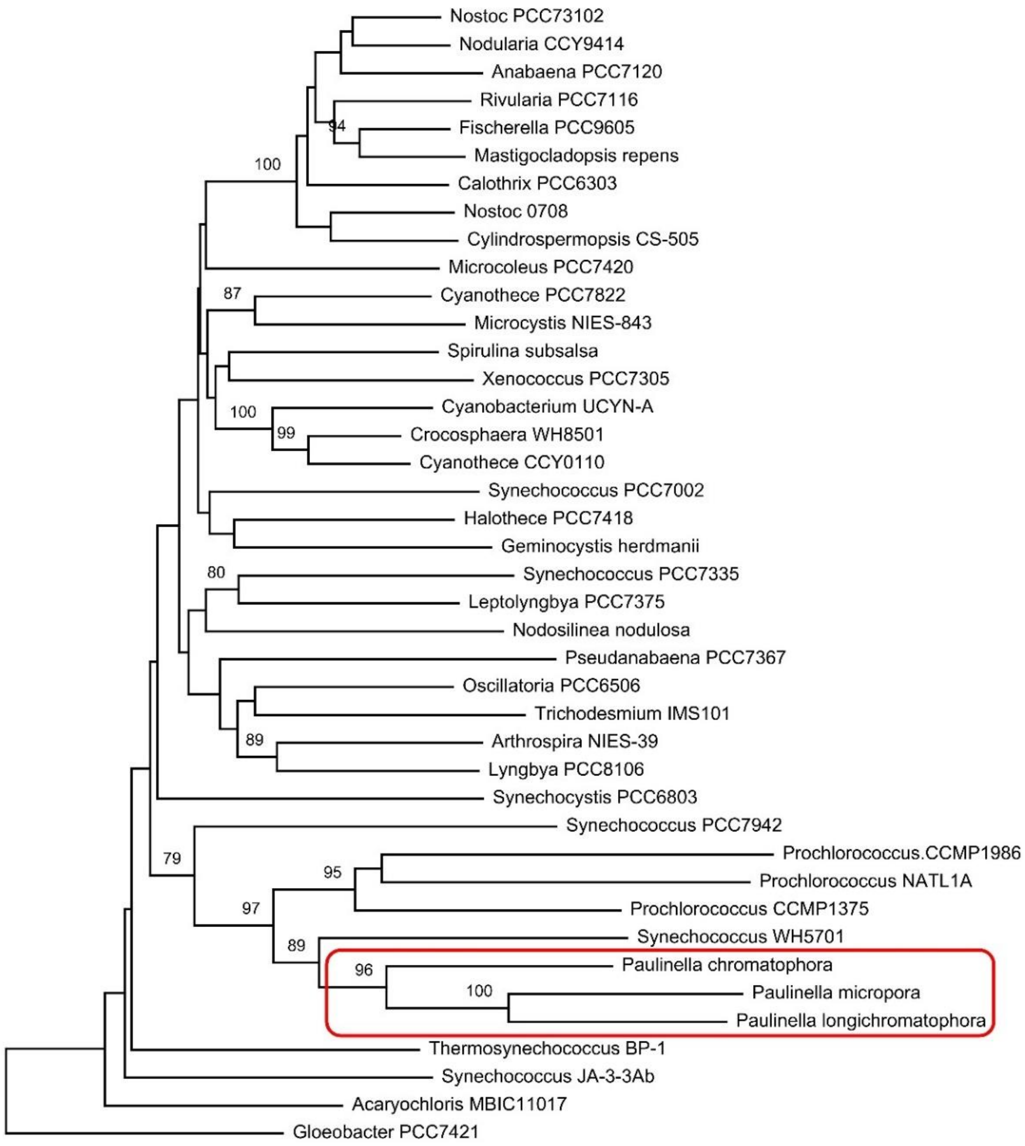


**Figure S1. Evolutionary relationships of RebA homologs.** The sequences of RebA homologs were retrieved from GenBank followed by a BLASTP search using RebA of Anabaena PCC 7120 as the query. The sequences from 38 cyanobacterial strains that could represent the diversity of cyanobacterial species (Shih et al., 2013) and 3 non-cyanobacterial strains of Paulinella were selected for phylogenetic analysis. These sequences were aligned and analyzed using MEGA X (Kumar et al., 2018). The evolutionary history was inferred using the Neighbor-Joining method (Saitou and Nei, 1987). Numbers above the branches are the percentage of replicate trees in which the associated taxa clustered together in the bootstrap test (500 replicates) (Felsenstein, 1985). Only the percentage values higher than 75% are shown.

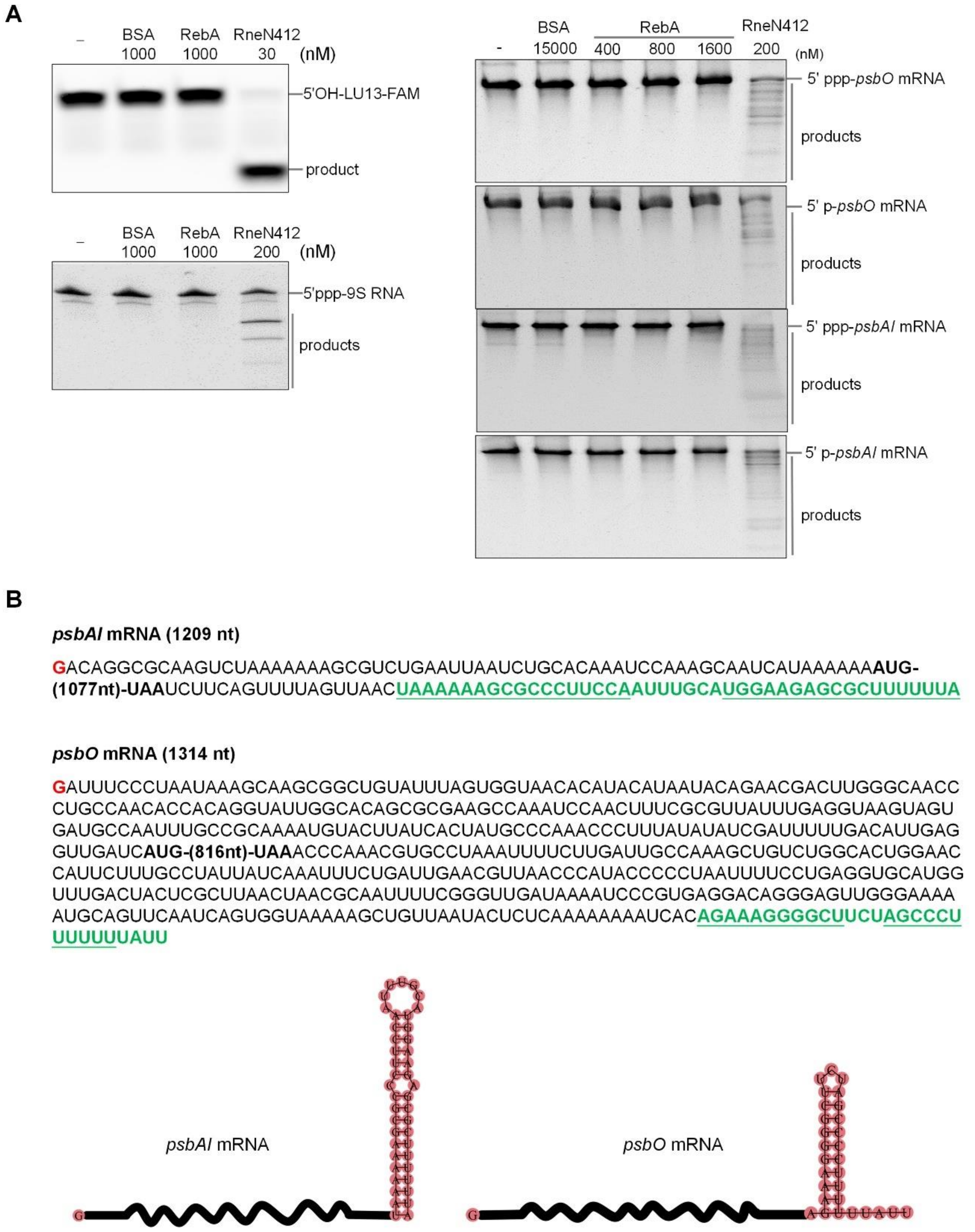


**Figure S2. Testing the RNA cleavage/degradation activities of RebA on various RNA substrates.** (A) RebA showed no cleavage/degradation activities on 5' OH-LU13-FAM, 5' ppp-9S RNA, *psbO* mRNA and *psbAI* mRNA (either with 5' p or 5' ppp). Each substrate was incubated with RebA in the reaction buffer, with BSA and RneN412 being used as a negative control and a positive control, respectively. In the reactions, 5' OH-LU13-FAM were used at 50 nM, 5' ppp-9S RNA, *psbO* mRNA and *psbAI* mRNAs were used at 500 nM. BSA, RebA and RneN412 were used at concentrations as indicated for each substrate. After 30 minutes of incubation, the reactions were stopped and analyzed by urea-PAGE. Substrates alone were loaded in the leftmost lanes (－) of the gels. (B) Sequence and cartoon representation of *psbA1* and *psbO* mRNAs used in this study. The initiation bases (in red) of these mRNAs were determined experimentally (Mitschke et al., 2011). The termination regions (in green) were predicted as they have features typical of a Rho-independent terminator (i.e., a stable stem-loop followed by a series of uridine residues). The termination residues that can form a stem by base-paring are underlined. The residues within the ORF regions are not shown for brevity.

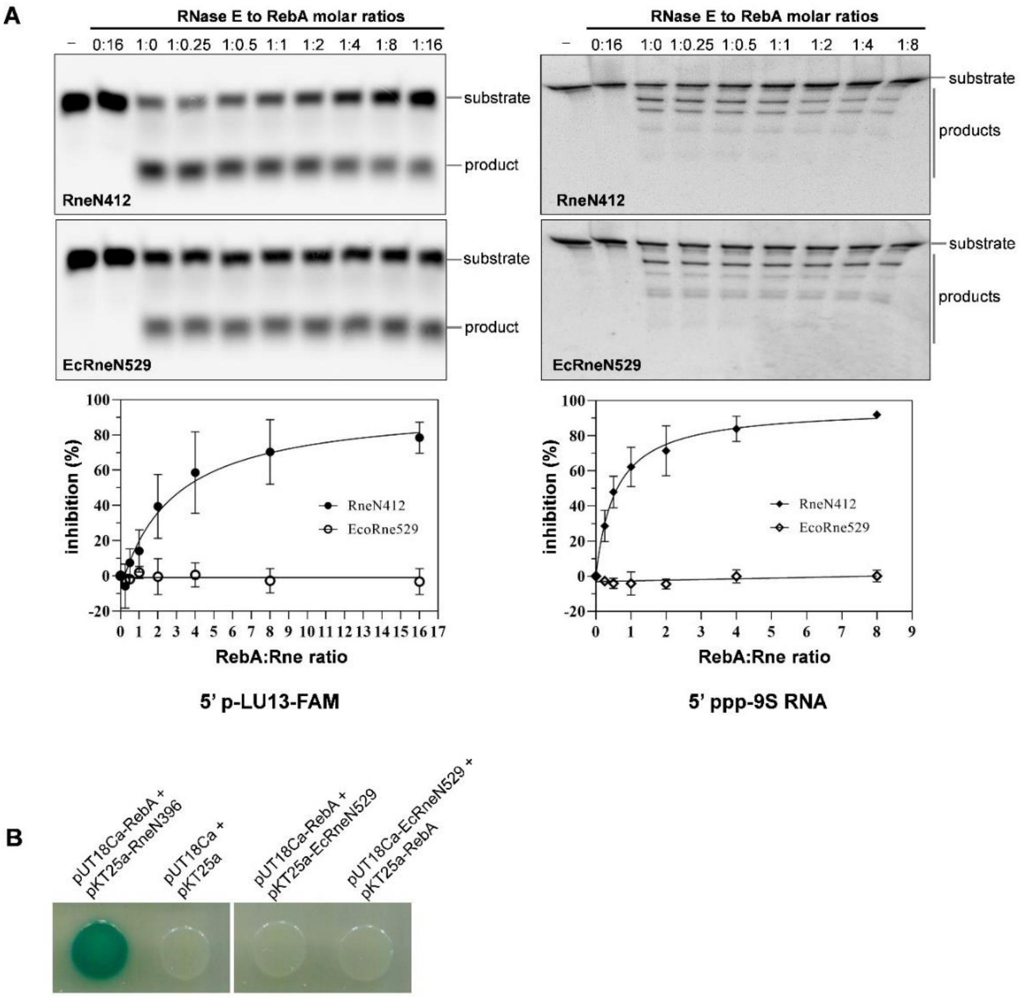


**Figure S3. Enzymatic and bacterial two hybrid assays showing that RebA neither inhibits the activities of nor interacts with *E. coli* RNase E.** (A) Comparing the inhibitory effect of RebA on the cleavage activity of the catalytic regions of *Anabaena* RNase E (RneN412) and *E. coli* RNase E (EcRneN529) using the substrate 5' p-LU13-FAM and 5' ppp-9S RNA. The assay conditions were identical to those used in Figure 2D. (B) Testing the interaction between RebA and EcRneN529 using bacterial two hybrid assay. The reporter strain containing the plasmids of pUT18Ca-RebA and pKT25a-RneN396 was used as a positive control and that containing the empty vectors of pUT18Ca and pKT25a was used as a negative control.

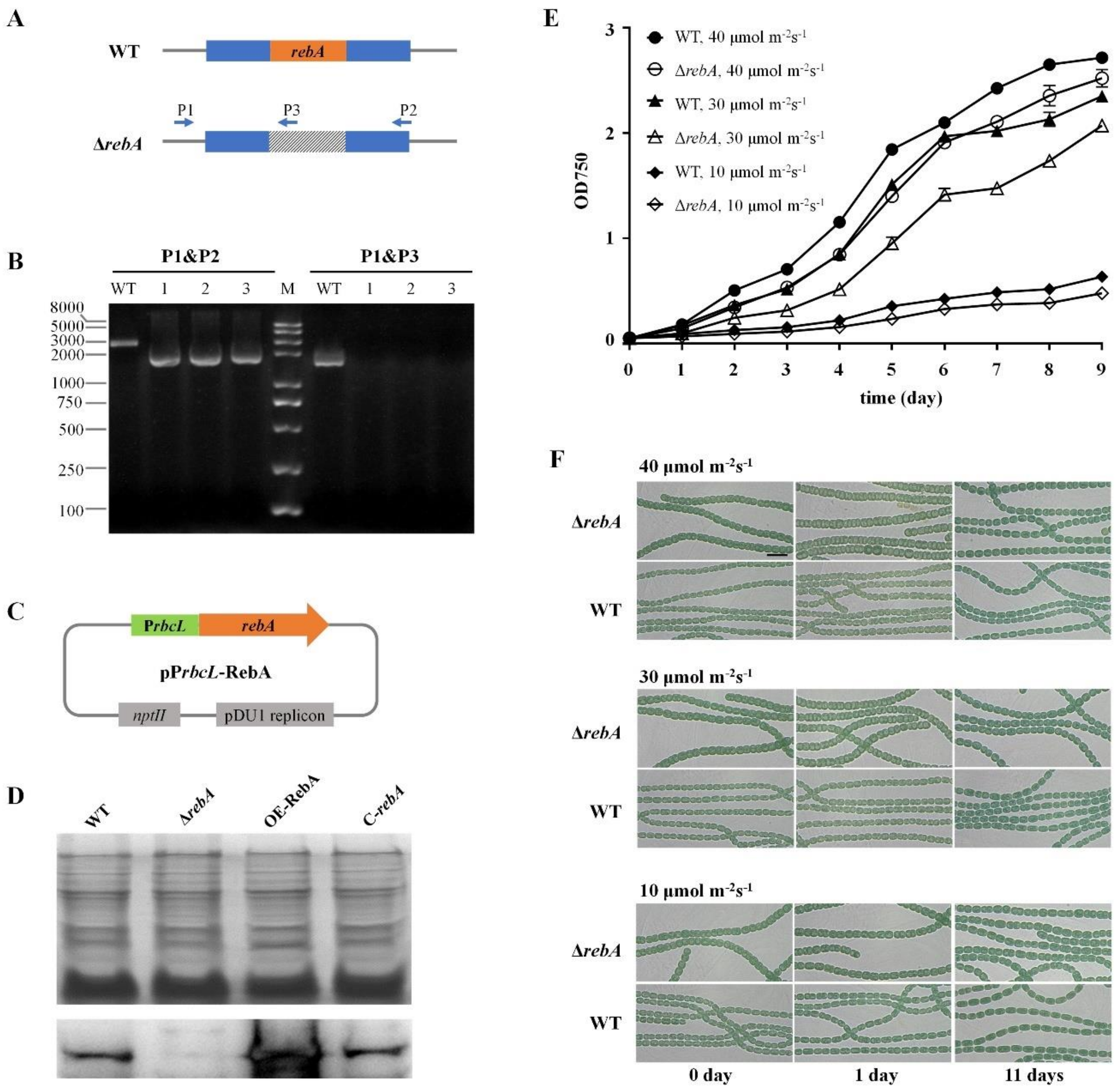


**Figure S4. Construction of *Anabaena rebA* mutation strain and overexpression strain, and checking the growth and cell morphology of the *rebA* mutant strain under different light intensities in BG11 medium.** (A) Schematic representation of the genotype of *Anabaena* wild type strain (WT) and the mutant strain of *rebA* (Δ*rebA*). (B) Verification on the genotype of Δ*rebA* by PCR using the primers shown in panel A (arrows). P1, P2, and P3 are short names for the primers of Pall1340F681, Pall1338R2225, and Pall1338R797, respectively (sequences listed in Table S3). Three colonies from the conjugation plates were examined. With P1 and P2, the expected sizes of the PCR products for WT and Δ*rebA* would be 3313 bp and 2086 bp, respectively. With P1 and P3, there would be a 1871 bp product for WT and no product for Δ*rebA*. (C) Schematic representation of the replicative plasmid pP*rbcL*-RebA, which was transferred into WT to obtain the RebA overexpression strain OE-RebA and into Δ*rebA* to obtain the complementation strain (C-*rebA*). Only the main features of the plasmid are shown. P*rbcL*, the promoter of the *Anabaena rbcL* gene that encodes the large-subunit of ribulose bisphosphate carboxylase; the promoter is a relative strong and often used for gene overexpression in *Anabaena*; *rebA*, the ORF of *rebA* gene; pDU1 replicon, the region required for plasmid replication in *Anabaena*; *nptII*, the neomycin resistance gene. (D) Comparing the expression of RebA in the strains of WT, Δ*rebA*, the overexpression strain OE-RebA and the complementary strain C-*rebA* by Western blot. The protein samples of the strains were loaded at 50 μg per lane onto duplicates of SDS-PAGE gels, with one gel stained with Coomassie brilliant blue staining (upper) and the other immunodetected with the polyclonal antibodies against RebA (lower). (E) Growth curves of Δ*rebA* and WT under high light (40 μmol $m^{-2}s^{-1}$), medium light (30 μmol $m^{-2}s^{-1}$) and low light (10 μmol $m^{-2}s^{-1}$) conditions. (F) Microscopic images of the Δ*rebA* and WT cells under the indicated light conditions. The shorter-cell phenotype of Δ*rebA* appeared when cell density was relatively low and light intensity was relatively high. The samples for microscopy were taken at the indicated time points during the process of growth curve measurement (panel E). Scale bar: 10 μm.

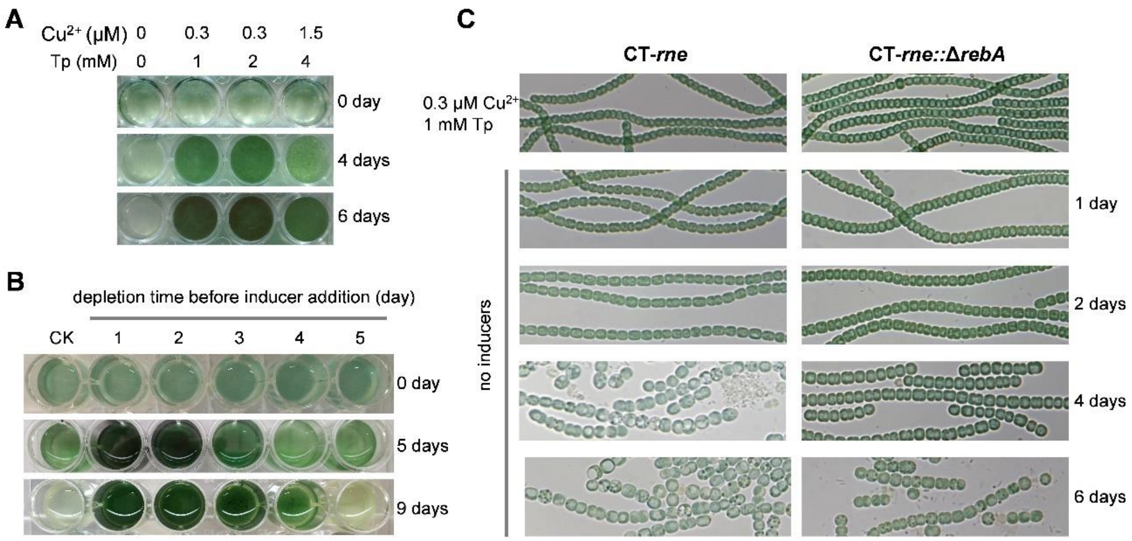


**Figure S5. Cell viability of the *rne* conditional mutant CT-*rne* under different induction conditions and the effects of *rebA* inactivation in CT-*rne*.** (A) Growth of CT-*rne* cells in the BG11 media with different concentrations inducers. (B) Viability of CT-*rne* cells after RNase E depletion in the medium without inducers. CT-*rne* cells precultured in the BG11 medium containing 0.3 μM $Cu^{2+}$ and 1 mM theophylline were inoculated into the wells of a 24-well plate containing the inducer-free medium with the initial density of OD700 ≈ 0.3. After indicated time periods of growth without inducers, 0.3 μM $Cu^{2+}$ and 1 mM theophylline were added into the wells. No inducers were added into the CK well. The cultures were grown in 24-well plates under normal condition (i.e., shaking speed of 180 rpm, light intensity of 30 μmol $m^{-2}s^{-1}$, and 30 °C). (C) The effects of *rebA* inactivation in CT-*rne*. The microscopic images of CT-*rne* and its derivate strain CT-*rne::* Δ*rebA*, in which *rebA* was additionally inactivated with the plasmid pCpf1b-ΔrebA, were compared. The strains were first cultured in BG11 medium containing 0.3 μM $Cu^{2+}$ and 1 mM theophylline (Tp), then transferred into the BG11 medium without the inducers for RNase E depletion. CT-*rne* cells became significantly longer after two days of RNase E depletion, and severe cell lysis and filament fragmentation occurred after four days of depletion. In contrast, when *rebA* was deleted in CT-*rne*, cell morphology change, filament fragmentation and cell lysis were significant delayed after RNase E depletion.

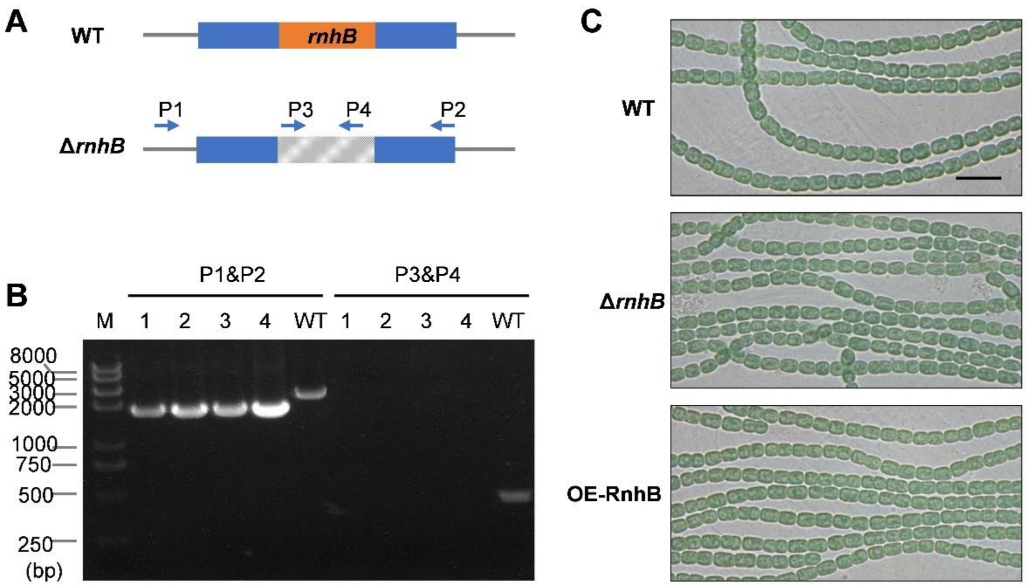


**Figure S6. Inactivation of *rnhB*, the gene downstream of *rne* did not affect cell morphology.** (A) Schematic representation of the genotype of *Anabaena* wild type strain (WT) and the mutant strain of *rnhB* (Δ*rnhB*). (B) Verification on the genotype of Δ*rnhB* by PCR using the primers shown in panel A (arrows). P1, P2, P3, and P4 are short names for the primers of Palr4332F928m, Palr4332R1538, Palr4332F179 and Palr4332R586, respectively (sequences listed in Table S3). Four colonies from the conjugation plates were examined. With P1 and P2, the expected sizes of the PCR products for WT and Δ*rnhB* would be 2466 bp and 1788 bp, respectively. With P3 and P4, there would be a 437 bp product for WT and no product for Δ*rnhB*. (C) Comparison of the microscopic images of the filaments of WT, Δ*rnhB* and the RnhB overexpression strain OE-RnhB. The samples for microscopy were taken from fresh cultures (OD700 ≈ 0.3) grown in BG11.

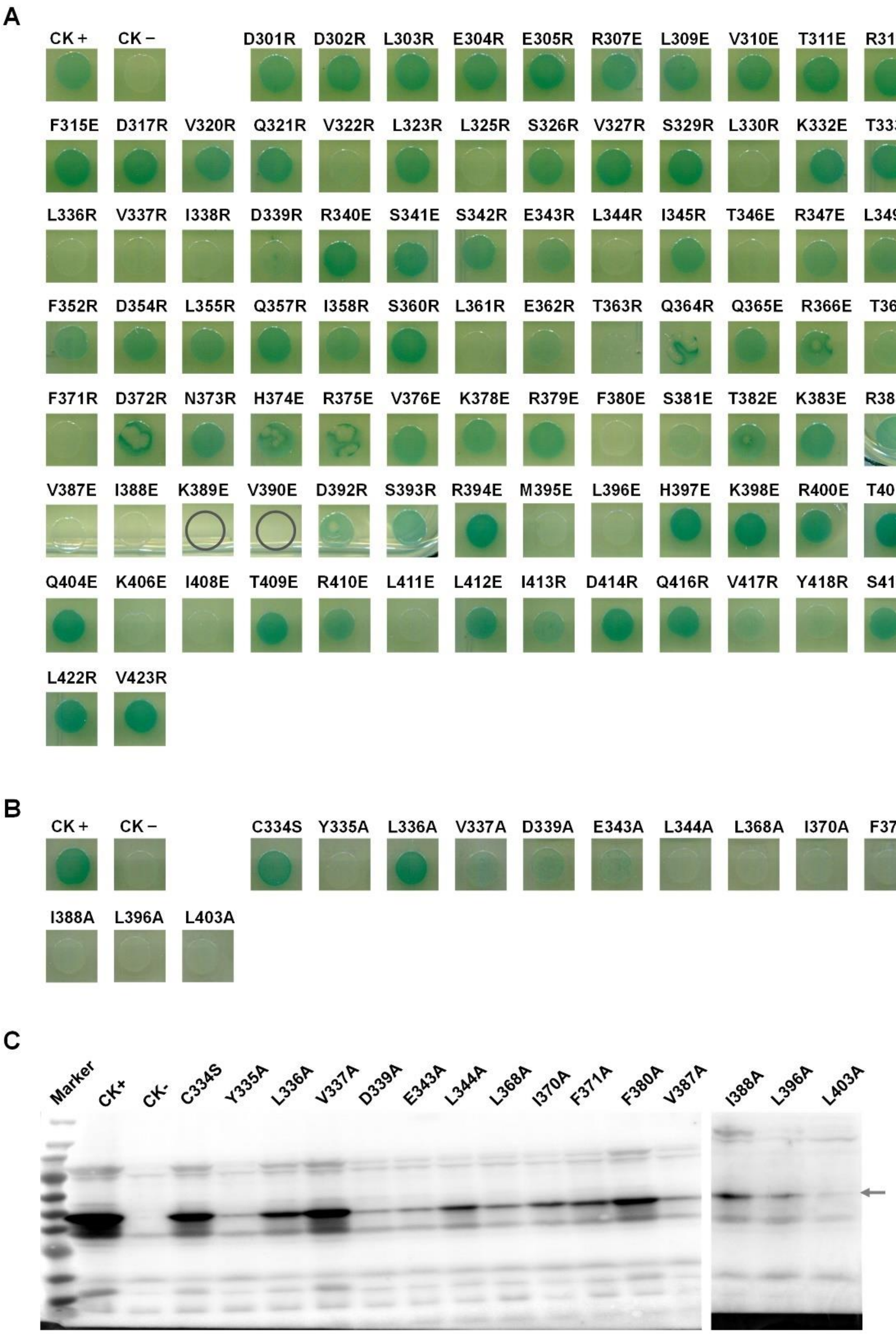


**Figure S7. Screening for the key residues required for RebA-RNase E interaction by site-directed mutagenesis and bacterial two hybrid assays.** Two rounds of screening were performed. (A) The first round of mutation screening. The residues of alanine, glycine and proline within RebA_C were not changed as they are unlikely involved interaction or their mutation may disrupt the protein structure. Creation of the mutations of K389E and V390E was not successful (indicated by circles). CK+ and CK-, the *E. coli* strains co-transformed with pUT18Ca-RneN396/pKT25a-RebA_C and pUT18C/pKT25, respectively. The other strains contained pUT18Ca-RneN396 and pKT25a-RebA_C derivates expressing the indicated RebA_C variants. (B) The second round of mutation screening. The sites whose mutation had showed significantly decreased interaction signal in the first round of screening were further changed into the small neutral residues of serine (for cysteine) or alanine (for other residues). (C) Confirming the expression of RebA_C variants in the two-hybrids strains used in panel B by Western blot. The antibodies against RebA were used. Arrow indicates the bands of RebA_C and its variants.

**Table S1. *Anabaena* and *E. coli* strains used in this study.** Abbreviations: Nm$^r$, neomycin resistance; Amp$^r$, ampicillin resistance; Cm$^r$, chloramphenicol resistance.

| Strains | Description | Source |
|---|---|---|
| *Anabaena* sp. PCC 7120 (WT) | the wild type strain used in this study | R. Haselkorn |
| OE-RebA | WT bearing the replicative plasmid pP$_{rbcL}$-RebA; Nm$^r$ | this study |
| Δ*rebA* | A markerless deletion mutant strain of *rebA* | this study |
| C-RebA | Δ*rebA* bearing pP$_{rbcL}$-RebA; Nm$^r$ | this study |
| CT-*rne* | *rne* conditional mutant strain, in which the native promoter of *rne* is replaced with the inducible CT promoter | this study |
| CT-*rne*::Δ*rebA* | CT-*rne* with additional deletion of *rebA* gene from the chromosome | this study |
| OE-RebA$^{V337A}$ | WT bearing the plasmid pP$_{rbcL}$-RebA$^{V337A}$; Nm$^r$ | this study |
| OE-RebA$^{I368A}$ | WT bearing pP$_{rbcL}$-RebA$^{I368A}$; Nm$^r$ | this study |
| OE-RebA$^{I370A}$ | WT bearing pP$_{rbcL}$-RebA$^{I370A}$; Nm$^r$ | this study |
| OE-RebA$^{V387A}$ | WT bearing pP$_{rbcL}$-RebA$^{V387A}$; Nm$^r$ | this study |
| OE-RebA$^{I388A}$ | WT bearing pP$_{rbcL}$-RebA$^{I388A}$; Nm$^r$ | this study |
| Δ*rnhB* | a markerless deletion mutant strain of *rnhB* | this study |
| OE-RnhB | WT bearing pCT-RnhB; Nm$^r$ | this study |
| BL21(DE3) | *E. coli* host strain for recombinant protein production | WEIDI Bio (CAT#: EC1002) |
| BTH101 | *E. coli* host strain (*cya*$^-$) for bacterial adenylate cyclase two-hybrid assay | Battesti and Bouveret, 2012 |
| DH10B (pRL443+pRL623) | *E. coli* strain used to transfer cargo plasmid into cyanobacteria by conjugation, containing the conjugal plasmid pRL443 and the helper plasmid pRL623; Amp$^r$, Cm$^r$. | Elhai et al., 1997 |

**Table S2. Plasmids used in this study.** Abbreviations: Km$^r$, kanamycin resistance; Nm$^r$, neomycin resistance; Sm$^r$, streptomycin resistance; Sp$^r$, spectinomycin resistance; Car$^r$, carbenicillin resistance.

| Plasmid | Description | Source |
|---|---|---|
| **for cyanobacteria strains construction** | | |
| pCT | Nm$^r$; a pDU1-derived vector that contains the inducible CT promoter | Zhou et al., 2020 |
| pCpf1b-sp | Sp$^r$/Sm$^r$; CRISPR/Cpf1-based genome editing vector | Niu et al., 2018 |
| pCpf1 | Nm$^r$; CRISPR/Cpf1-based genome editing vector | Niu et al., 2018 |
| pICT-rne | Sp$^r$/Sm$^r$; the genome editing plasmid for *rne* conditional mutation | this study |
| pCpf1b-ΔrebA | Sp$^r$/Sm$^r$; the genome editing plasmid for *rebA* markerless deletion | this study |
| pP$_{rbcL}$-RebA | Nm$^r$; pRL25N-derived plasmid for RebA expression under the relatively strong promoter P$_{rbcL}$. | this study |
| pP$_{rbcL}$-RebA$^{V337A}$ | Nm$^r$; pP$_{rbcL}$-RebA derivate for expressing the RebA variant that has the V337A mutation | this study |
| pP$_{rbcL}$-RebA$^{I368A}$ | Nm$^r$; pP$_{rbcL}$-RebA derivate for expressing the RebA variant that has the I368A mutation | this study |
| pP$_{rbcL}$-RebA$^{I370A}$ | Nm$^r$; pP$_{rbcL}$-RebA derivate for expressing the RebA variant that has the I370A mutation | this study |
| pP$_{rbcL}$-RebA$^{V387A}$ | Nm$^r$; pP$_{rbcL}$-RebA derivate for expressing the RebA variant that has the V387A mutation | this study |
| pP$_{rbcL}$-RebA$^{I388A}$ | Nm$^r$; pP$_{rbcL}$-RebA derivate for expressing the RebA variant that has the I388A mutation | this study |
| pCpf1b-ΔrnhB | Sp$^r$/Sm$^r$; the genome editing plasmid for *rnhB* markerless deletion | this study |
| pCT-RnhB | Nm$^r$; pCT carrying *rnhB* gene under the control of the CT promoter, used for RnhB overexpression | this study |
| **for recombinant protein expression** | | |
| pET28a | Km$^r$; vector for expressing recombinant protein with an N-terminal His tag and/or a C-terminal His tag. | Invitrogen |
| pHStag | Km$^r$; pET28a-derived vector for expressing recombinant protein with an C-terminal Strep tag and/or a N-terminal His tag. | Zhou et al., 2020 |
| pNStrep | Km$^r$; pET28a-derived vector for expressing recombinant protein with an N-terminal Strep tag and/or a C-terminal His tag. | this study |
| pHTAlr4331N412 | Km$^r$; for expressing the catalytic domain of *Anabaena* RNase E (1–412 aa) with an N-terminal His tag | Zhang et al., 2014 |
| pHTAlr4331N396 | Km$^r$; for expressing the catalytic domain of *Anabaena* RNase E (1–396 aa) with an N-terminal His tag | this study |

| | | |
|---|---|---|
| pHT-EcRneN529 | $Km^r$; for expressing the catalytic domain of *E. coli* RNase E (1–529 aa) with an N-terminal His tag | this study |
| pNStrep-RebA | $Km^r$; for expressing RebA with an N-terminal Strep tag; the ORF of rebA was codon-optimized and cloned into the vector pNStrep. | this study |
| pNStrep-RebA$^{V337A}$ | $Km^r$; same as pNStrep-RebA, but for expressing the RebA variant having the point mutation V337A | this study |
| pNStrep-RebA$^{L368A}$ | $Km^r$; same as pNStrep-RebA, but for expressing the RebA variant having the point mutation I368A | this study |
| pNStrep-RebA$^{I370A}$ | $Km^r$; same as pNStrep-RebA, but for expressing the RebA variant having the point mutation I370A | this study |
| pNStrep-RebA$^{V387A}$ | $Km^r$; same as pNStrep-RebA, but for expressing the RebA variant having the point mutation V387A | this study |
| pNStrep-RebA$^{I388A}$ | $Km^r$; same as pNStrep-RebA, but for expressing the RebA variant having the point mutation I388A | this study |
| **for bacterial two-hybrid assay** | | |
| pKT25a | $Km^r$; vector for two-hybrid assay, encoding the adenylate cyclase fragment T25. Derived from pKT25 (Battesti and Bouveret, 2012) by changing the cloning site for easier cloning. | this study |
| pUT18Ca | $Car^r$; vector for two-hybrid assay, encoding the adenylate cyclase fragment T18. Derived from pUT18C (Battesti and Bouveret, 2012) by changing the cloning site of pUT18 to that of pKT25a, so that a same fragment can be cloned into both pKT25a and pUT18Ca. | this study |
| pKT25a-zip | $Km^r$; positive control plasmid used in bacterial two-hybrid assay | Battesti and Bouveret, 2012 |
| pUT18C-zip | $Car^r$; positive control plasmid used in bacterial two-hybrid assay | Battesti and Bouveret, 2012 |
| pKT25a-RebA | $Km^r$; plasmid derived from pKT25, encoding RebA fused to the C-terminal of T25. | this study |
| pUT18Ca-RebA | $Car^r$; pUT18Ca derivate encoding RebA fused to the C-terminal of T18. | this study |
| pKT25a-RebA_C | $Km^r$; pKT25 derivate encoding RebA_C (301-424 aa) fused to the C-terminal of T25. | this study |
| pUT18Ca-RebA_C | $Car^r$; pUT18Ca derivate encoding RebA_C (301-424 aa) fused to the C-terminal of T18. | this study |
| pKT25a-Alr4331 | $Km^r$; pKT25 derivate encoding RNase E fused to the C-terminal of T25. | Zhou et al., 2020 |
| pUT18Ca-Alr4331 | $Car^r$; pUT18Ca derivate encoding RNase E fused to the C-terminal of T18. | Zhou et al., 2020 |
| pKT25a-Alr4331N | $Km^r$; pKT25 derivate encoding RNase E N-terminal region (1-396 aa, the catalytic region) fused to the C-terminal of T25. | Zhou et al., 2020 |
| pUT18Ca-Alr4331N | $Car^r$; pUT18Ca derivate encoding RNase E N-terminal region (1-396 aa, the catalytic region) fused to the C-terminal of T18. | Zhou et al., 2020 |
| pKT25a-Alr4331(1-117) | $Km^r$; pKT25 derivate encoding RNase E N-terminal region (1-117 aa, the S1 domain) fused to the C-terminal of T25. | this study |

| | | |
|---|---|---|
| pUT18Ca-Alr4331(1-117) | Car$^r$; pUT18Ca derivate encoding RNase E N-terminal region (1-117 aa, the S1 domain) fused to the C-terminal of T18. | this study |
| pKT25a-Alr4331(100-210) | Km$^r$; pKT25 derivate encoding RNase E N-terminal region (100-210 aa, the 5’ sensor domain) fused to the C-terminal of T25. | this study |
| pUT18Ca-Alr4331(100-210) | Car$^r$; pUT18Ca derivate encoding RNase E N-terminal region (100-210 aa, the 5’ sensor domain) fused to the C-terminal of T18. | this study |
| pKT25a-Alr4331(196-273) | Km$^r$; pKT25 derivate encoding RNase E N-terminal region (196-273 aa, the RNase H domain) fused to the C-terminal of T25. | this study |
| pUT18Ca-Alr4331(196-273) | Car$^r$; pUT18Ca derivate encoding RNase E N-terminal region (196-273 aa, the RNase H domain) fused to the C-terminal of T18. | this study |
| pKT25a-Alr4331(255-396) | Km$^r$; pKT25 derivate encoding RNase E N-terminal region (255-396 aa, the DNase I domain) fused to the C-terminal of T25. | this study |
| pUT18Ca-Alr4331(255-396) | Car$^r$; pUT18Ca derivate encoding RNase E N-terminal region (225-396 aa, the DNase I domain) fused to the C-terminal of T18. | this study |
| pKT25a-All4396 | Km$^r$; pKT25 derivate encoding PNPase fused to the C-terminal of T25. | this study |
| pUT18Ca-All4396 | Car$^r$; pUT18Ca derivate encoding PNPase fused to the C-terminal of T18. | this study |
| pKT25a-Alr1240 | Km$^r$; pKT25 derivate encoding RNase II fused to the C-terminal of T25. | Zhou et al., 2020 |
| pUT18Ca-Alr1240 | Car$^r$; pUT18Ca derivate encoding RNase II fused to the C-terminal of T18. | Zhou et al., 2020 |
| pKT25a-Alr1223 | Km$^r$; pKT25 derivate encoding CrhB fused to the C-terminal of T25. | this study |
| pUT18Ca-Alr1223 | Car$^r$; pUT18Ca derivate encoding CrhB fused to the C-terminal of T18. | this study |
| PpKT25a-EcRneN529 | Km$^r$; pKT25 derivate encoding *E. coli* RNase E N-terminal region (1-529 aa) fused to the C-terminal of T25. | this study |
| pUT18Ca-EcRneN529 | Car$^r$; pUT18Ca derivate encoding *E. coli* RNase E N-terminal region (1-529 aa) to the C-terminal of T18. | this study |

**Table S3. Oligonucleotides used in this study. Related to STAR Methods.**

| Name | Sequence (5'-3') | Description |
|---|---|---|
| **for construction of cyanobacterial plasmids** | | |
| Pall1338F1005m | GCAGAAATTCGATATCTAGATCGTACTGAGTCCTGAGGTAGAAGTCA | to amplify a region upstream *rebA* ORF, which is used to construct the plasmid pCpf1b-ΔrebA |
| Pall1338R11 | AGCGCTACCGACGCTAGTGCTTTGTTAGTTTTCTCACGGTAGATAC | |
| Pall1338R2225 | CGCAACGTTGTTGCCATTGCAGGAACTTGGTCTGATGACAG | to amplify a region downstream *rebA* ORF, which is used to construct the plasmid pCpf1b-ΔrebA |
| Pall1338F1247a | GCACTAGCGTCGGTAGCGCTCAGGTCTATTCTTTGTCTCTGGTGTA | |
| cr_all1338R1130F | AGATGCCACACGGTGGTTATCAAATA | to generate the DNA duplex of a spacer sequence, which is used to construct the plasmid pCpf1b-ΔrebA |
| cr_all1338R1130R | AGACTATTTGATAACCACCGTGTGGC | |
| Pall1340F681 | TGAGGAGTTATTTGCTAGTGCCAGTAACTATTGGCAGTTGCGAG | to verify the genotype of Δ*rebA* strain |
| Pall1338R2225 | CGCAACGTTGTTGCCATTGCAGGAACTTGGTCTGATGACAG | |
| Pall1338R797 | GGTCTTGCGATTACCTGCGGTGGAGCA | |
| P25TNotI-F | AAGCCCAAGCGTTTTGTTATTGG | to amplify the vector fragment from pCT, which is used to construct the plasmid pPrbcL-RebA |
| P25TBamHI-R | GGATCCAAAAAAAAACCCCGCCGAAGC | |
| PPrbcL-F | CGGGGTTTTTTTTTGGATCCGCAGGGGAAGTAAAGAAGAATGA | to amplify the promoter of PrbcL from *Anabaena* genomic DNA, which is used to construct the plasmid pPrbcL-RebA |
| PPrbcL-R | CTCGAGTATCCTTCCAAGATGTC | |
| Pall1338bF1c | CTTGGAAGGATACTCGAGATGCGCAAACTGACCAACAGCGACAA | to amplify the *rebA* gene from pNStrep-RebA, which is used to construct the plasmid pPrbcL-RebA |
| Pall1338bR1272c | AACAAAACGCTTGGGCTTTTACACCAGGCTCAAGGAGTAGACTT | |
| PAll1338bV337A_f | ACCTGGCCATTGATCGTAGCTCCGAACTG | to generate the mutation V337A in RebA by site-directed mutagenesis, using the template of pPrbcL-RebA or pNStrep-RebA. |
| PAll1338bV337A_r | TCAATGGCCAGGTAGCAGGTTTTCGGCA | |
| PAll1338bL368A_f | CCGCGCCGATTTTCGACAACCACCGT | same as above, but to generate the mutation L368A in RebA. |
| PAll1338bL368A_r | AAAATCGGCGCGGTGCGCTGCTGGGT | |
| PAll1338bI370A_f | GCCGGCGTTCGACAACCACCGTGTG | same as above, but to generate the mutation I370A in RebA. |
| PAll1338bI370A_r | GTCGAACGCCGGCAGGGTGCGCTG | |
| PAll1338bV387A_f | GCGCAATTAAGGTTCCGGATAGCCGTA | same as above, but to generate the mutation V387A in RebA. |
| PAll1338bV387A_r | ACCTTAATTGCGCGGTCACGCTTGGTG | |
| PAll1338bI388A_f | GTTGCGAAGGTTCCGGATAGCCGTAT | |

| | | |
|---|---|---|
| PAll1338bI388A_r | GAACCTTCGCAACGCGGTCACGCTTG | same as above, but to generate the mutation I337A in RebA. |
| Palr4331F1596m | GCAGAAATTCGATATCTAGATGCGATCGCCCAAGCCGGTCTA | to amplify a region upstream *rne* ORF, which is used to construct the plasmid pICT-rne. |
| Palr4331R556m | GGCTCGACTCTAGCTAGAGTTATACAGCTAAAATTAGTTGATTGCGA | |
| PcoquwF | CTCTAGCTAGAGTCGAGCCCGTTAAGGGATTTTGGTCATGAG | to amplify the CT promoter from the plasmid pCT, which is used to construct the plasmid pICT-rne |
| PV_19 | CATCTTGTTGTTACCTCCTTAGCA | |
| Palr4331F1b | TGCTAAGGAGGTAACAACAAGATGCCAAAACAAATTATTATCGCGG | to amplify a 5' region of *rne* ORF, which is used to construct the plasmid pICT-*rne* |
| Palr4331R990 | CGCAACGTTGTTGCCATTGCCCGCAGACGTAGCTGGCGAGCA | |
| cr_alr4331F26mF | AGATATACTGCCGATTTTTGAGGAAA | to generate the DNA duplex of a spacer sequence, which is used to construct the plasmid pICT-*rne*. |
| cr_alr4331F26mR | AGACTTTCCTCAAAAATCGGCAGTAT | |
| Pall0258F475m | CTTGGATCCTAAAGCCTGTGAAATTAACTG | to verify the genotype of the strain CT-*rne*. |
| Palr4331R1020 | GAAATCAACGACGATTACCCCAGCGAT | |
| Palr4331F1596m | GCAGAAATTCGATATCTAGATGCGATCGCCCAAGCCGGTCTA | |
| Palr4331R33m | AAGGGCATCAAGACGATGCTGGTATCACCGCTTGGAGGTTTGTTATCCCT | |
| Pall3854F219ma | ATCCTAATACGACTCACTATAGATTTCCCTAATAAAGCAAGCGGCTGTAT | to amplify the DNA template for *in vitro* transcription of *psbO* mRNA by T7 RNA polymerase |
| Pall3854R1095 | AATAAAAAAAGGGCTAGAAGCCCCTTTCTGTGATT | |
| Palr4866F65maa | ATCCTAATACGACTCACTATAGACAGGCGCAAGTCTAAAAAAAGCGTCTGAATTA | to amplify the DNA template for in vitro transcription of *psbAI* mRNA by T7 RNA polymerase |
| Palr4866R1144a | TAAAAAAGCGCTCTTCCATGCAAATTGGAAGGGCGCTTTTTTAGTTAACTAAAACTGA | |
| Palr4332F797m | TGGCAGAAATTCGATATCTAGATCAGTGAGTCAGATTTAGACCTTGATGG | to amplify a region upstream *rnhB* ORF, which is used to construct the plasmid pCpf1b-ΔrnhB. |
| Palr4332R1m | CAAAGCTTACACTAATACACAATATCTCA | |
| Palr4332F679 | TGTATTAGTGTAAGCTTTGTTAATTCGTAATTAAACTGTTCTTTTCTTTCAACA | to amplify a region downstream *rnhB* ORF, which is used to construct the plasmid pCpf1b-ΔrnhB. |
| Palr4332R1464 | CGCAACGTTGTTGCCATTGCAACTAAGTGGCATTTAGCCCATCAACTA | |
| cr_alr4332R379F | AGATGCTTTTTTACAGACCGTTTCAT | to generate the DNA duplex of a spacer sequence, which is used to construct the plasmid pCpf1b-ΔrnhB. |
| cr_alr4332R379R | AGACATGAAACGGTCTGTAAAAAAGC | |
| Palr4332F928m | TAATCATCCTAGCTATCAAGAACTGAAC | to verify the genotype of the strain Δ*rnhB.* |
| Palr4332R586 | CATAGCCCTTGTTACTCTTTAAGTCATACA | |
| Palr4332F179 | GCCTGTAGTGGCAGCATCTGTCATATTACC | |

| | | |
|---|---|---|
| Palr4332R1538 | ATGAGCTATAATCTATATACTTACGAACCA | |
| **for construction of protein expression plasmids** | | |
| PV_1 | GCTGCCGCGCGGCACCAG | to amplify the vector fragment from pET28a, which is used to construct the plasmid pHT-EcRneN529. |
| PV_2 | CTGAGATCCGGCTGCTAAC | |
| Pb1084F1 | CTGGTGCCGCGCGGCAGCATGAAAAGAATGTTAATCAACGCAACTCAGCAGGAAGAG | to amplify a region of *E. coli rne* gene, which is used to construct the plasmid pHT-EcRneN529. |
| Pb1084R1587 | GTTAGCAGCCGGATCTCAGTTACAGCGCAGGTTGTTCCGGACGCTTACGTTCA | |
| PV_5 | TGCACCCTTTTCGAACTGC | to amplify the vector fragment from pNStrep, which is used to construct the plasmids pNStrep-Alr4331N396 and pNStrep-RebA. |
| PV_2 | CTGAGATCCGGCTGCTAAC | |
| Palr4331F4b | CAGTTCGAAAAGGGTGCAATGCCAAAACAAATTATTATCGCGGAGCA | to construct the plasmid pNStrep-Alr4331N396. |
| Palr4331R1188b | GTTAGCAGCCGGATCTCAGTTAAGTATCACCAAACAATTCATAAATATTTTGCCCT | |
| Pall1338F901 | AACAGGTACCCATATGGACGACTTAGAGGAGGCAAGACC | to amplify a *rebA* region from Anabaena chromosome, which is used to construct the plasmid pHS-RebA_C. |
| Pall1338R1269 | TTCTACTCGAGTCACCAGAGACAAAGAATAGACCT | |
| Pall1338F2 | AACAGGTACCCATATGAGAAAACTAACAAACTCTG | to amplify a *rebA* region from Anabaena chromosome, which is used to construct the plasmid pHS-RebA_DC. |
| Pall1338R960 | AATCTCGAGTCACTGCGGCATCCCCAAATC | |
| **for construction of bacterial two-hybrid assay plasmids** | | |
| PpKT25a-F733 | GAGATCTAGATCGACATCTGTCGAGC | to prepare the linear fragment of the vector pKT25a or pUT18C. |
| PpKT25a-R720 | TTCACCACTAGAGGTGATCA | |
| Pall1338F1b | ATCACCTCTAGTGGTGAAGTGAGAAAACTAACAAACTCTGACAA | to amplify the *Anabaena* genomic region encoding RebA (2-423 aa) |
| Pall1338R1272a | GATGTCGATCTAGATCTCTTACACCAGAGACAAAGAATAGACCT | |
| Pall1338bF4 | ATCACCTCTAGTGGTGAACGCAAACTGACCAACAGCGACAAGCAGGA | to amplify the *Anabaena* genomic region encoding RebA (2-423 aa). |
| Pall1338bR1272b | GATGTCGATCTAGATCTCTTACACCAGGCTCAAGGAGTAGACTT | |
| Palr4331F1c | ATCACCTCTAGTGGTGAAATGCCAAAACAAATTATTATCGCGG | to amplify the *Anabaena* genomic region encoding RNase E (1-688 aa). |
| Palr4332R2094 | GATGTCGATCTAGATCTCAATCTGTGGCTCTGCTTCG | |
| Palr4331F4 | ATCACCTCTAGTGGTGAACCAAAACAAATTATTATCGCGGAGCAGCATC | to amplify the *Anabaena* genomic region encoding RNase E N terminal domain (2-396 aa). |
| Palr4331R1188 | GATGTCGATCTAGATCTCTTAAGTATCACCAAACAATTCATAAATATTTTGCCCT | |
| PpKT25a-F733 | GAGATCTAGATCGACATCTGTCGAGC | |

| | | |
|---|---|---|
| Palr4331R351 | GTCGATCTAGATCTCTTAACGTCCAGGCAGAGTGATATTACCTGTGAG | to amplify pKT25a-Alr4331N or pUT18Ca-Alr4331N for constructing pKT25a-Alr4331 (2-117) or pUT18Ca-Alr4331 (2-117) |
| Palr4331F298 | ATCACCTCTAGTGGTGAACCAACGGGAACAAAAGGCCCAAGGCTCACA | to amplify the *Anabaena* genomic region encoding RNase E N terminal domain (100-210 aa). |
| Palr4331R630 | GATGTCGATCTAGATCTCACGTAGTACGCGCTGGATAAAGTCAT | |
| Palr4331F586 | ATCACCTCTAGTGGTGAAGCACTCCTGAATCGGGACGATGACTTTATCC | to amplify the *Anabaena* genomic region encoding RNase E N terminal domain (196-273 aa). |
| Palr4331R819 | GATGTCGATCTAGATCTCTTTAAGGGCTTCTCGAATTGCGGCATTGA | |
| Pall1338F901 | ATCACCTCTAGTGGTGAAGACGACTTAGAGGAGGCAAGACCTCTAGTTAC | to amplify the *Anabaena* genomic region encoding RebA C terminal domain (300-424 aa). |
| Pall1338R1272a | GATGTCGATCTAGATCTCTTACACCAGAGACAAAGAATAGACCT | |
| Palr4331F763 | ACCTCTAGTGGTGAACGCTCCCCAATTTTAGAATATTTCC | to amplify pKT25a-Alr4331N or pUT18Ca-Alr4331N for constructing pKT25a-Alr4331 (255-396 aa) or pUT18Ca-Alr4331 (255-396 aa) |
| PpKT25a-R720 | TTCACCACTAGAGGTGATCA | |
| PAll1338_C334S_f | ACCTCATATTTGGTTATTGATCGCTCCTCG | to create the C334S mutation in RebA in the two-hybrid plasmids of RebA (i.e., pKT25-RebA and pUT18Ca-RebA). |
| PAll1338_C334S_r | CCAAATATGAGGTCTTGGGCAAAGATGCC | |
| PAll1338_Y335A_f | CTGCGCATTGGTTATTGATCGCTCCTCG | same as above, but for the mutation Y335A |
| PAll1338_Y335A_r | AACCAATGCGCAGGTCTTGGGCAAAGAT | |
| PAll1338_L336A_f | GCTATGCAGTTATTGATCGCTCCTCGGAG | same as above, but for the mutation L336A |
| PAll1338_L336A_r | ATAACTGCATAGCAGGTCTTGGGCAAAGA | |
| PAll1338_V337A_f | ATTTGGCCATTGATCGCTCCTCGGAGTTA | same as above, but for the mutation V337A |
| PAll1338_V337A_r | TCAATGGCCAAATAGCAGGTCTTGGGCAA | |
| PAll1338_I338A_f | GGTTGCAGATCGCTCCTCGGAGTTAATTAC | same as above, but for the mutation I338A |
| PAll1338_I338A_r | GCGATCTGCAACCAAATAGCAGGTCTTGGG | |
| PAll1338_D339A_f | GGTTATTGCACGCTCCTCGGAGTTAATTACC | same as above, but for the mutation D339A |
| PAll1338_D339A_r | GCGTGCAATAACCAAATAGCAGGTCTTGGG | |
| PAll1338_E343A_f | TCGGCTTTAATTACCAGACCACTCAAAGACTT | same as above, but for the mutation E343A |
| PAll1338_E343A_r | TAATTAAAGCCGAGGAGCGATCAATAACCAAA | |
| PAll1338_L344A_f | GAGGCCATTACCAGACCACTCAAAGACTTTG | same as above, but for the mutation L344A |
| PAll1338_L344A_r | TGGTAATGGCCTCCGAGGAGCGATCAATAAC | |
| PAll1338_T346A_f | TAATTGCAAGACCACTCAAAGACTTTGGTG | same as above, but for the mutation T346A |
| PAll1338_T346A_r | GGTCTTGCAATTAACTCCGAGGAGCGATCA | |
| PAll1338_R347A_f | TACCGCTCCACTCAAAGACTTTGGTGATTTG | same as above, but for the mutation R347A |

| | | |
|---|---|---|
| PAll1338_R347A_r | GAGTGGAGCGGTAATTAACTCCGAGGAGCGA | |
| PAll1338_L349A_f | AGACCAGCGAAAGACTTTGGTGATTTGGGGC | same as above, but for the mutation L349A |
| PAll1338_L349A_r | CTTTCGCTGGTCTGGTAATTAACTCCGAGG | |
| PAll1338_D351A_f | CTCAAAGCCTTTGGTGATTTGGGGCAAATTCC | same as above, but for the mutation D351A |
| PAll1338_D351A_r | CAAAGGCTTTGAGTGGTCTGGTAATTAACTCC | |
| PAll1338_F352A_f | CAAAGACGCCGGTGATTTGGGGCAAATTCCTA | same as above, but for the mutation F352A |
| PAll1338_F352A_r | ACCGGCGTCTTTGAGTGGTCTGGTAATTAACT | |
| PAll1338_L361A_f | CCTAGTGCAGAAACCCAGCAAAGAACACTAC | same as above, but for the mutation L361A |
| PAll1338_L361A_r | TTTCTGCACTAGGAATTTGCCCCAAATCAC | |
| PAll1338_E362A_f | AGTCTAGCCACCCAGCAAAGAACACTACCA | same as above, but for the mutation E362A |
| PAll1338_E362A_r | GGGTGGCTAGACTAGGAATTTGCCCCAAAT | |
| PAll1338_T363A_f | GAAGCGCAGCAAAGAACACTACCAATATTTGAT | same as above, but for the mutation T363A |
| PAll1338_T363A_r | TTTGCTGCGCTTCTAGACTAGGAATTTGCCCCA | |
| PAll1338_T367A_f | AAAGAGCACTACCAATATTTGATAACCACCGTG | same as above, but for the mutation T367A |
| PAll1338_T367A_r | GGTAGTGCTCTTTGCTGGGTTTCTAGACTAGG | |
| PAll1338_L368A_f | GAACAGCGCCAATATTTGATAACCACCGTGTG | same as above, but for the mutation L368A |
| PAll1338_L368A_r | ATTGGCGCTGTTCTTTGCTGGGTTTCTAGAC | |
| PAll1338_I370A_f | CTACCAGCTTTTGATAACCACCGTGTGGC | same as above, but for the mutation I370A |
| PAll1338_I370A_r | CAAAAGCTGGTAGTGTTCTTTGCTGGGT | |
| PAll1338_F371A_f | CCAATAGCTGATAACCACCGTGTGGCTAAA | same as above, but for the mutation F371A |
| PAll1338_F371A_r | TATCAGCTATTGGTAGTGTTCTTTGCTGGG | |
| PAll1338_F380A_f | ACGCGCGTCTACCAAGCGCGATCGCGT | same as above, but for the mutation F380A |
| PAll1338_F380A_r | GGTAGACGCGCGTTTAGCCACACGGT | |
| PAll1338_S381A_f | CGCTTTGCCACCAAGCGCGATCGCGT | same as above, but for the mutation S381A |
| PAll1338_S381A_r | TGGTGGCAAAGCGTTTAGCCACACGG | |
| PAll1338_D385A_f | AGCGCGCACGCGTGATTAAAGTCCCCG | same as above, but for the mutation D385A |
| PAll1338_D385A_r | ACGCGTGCGCGCTTGGTAGAAAAGCGT | |
| PAll1338_R386A_f | ATGCTGTGATTAAAGTCCCCGATAGTAGAAT | same as above, but for the mutation R386A |
| PAll1338_R386A_r | TTAATCACAGCATCGCGCTTGGTAGAAAAGC | |
| PAll1338_V387A_f | CGCTATTAAAGTCCCCGATAGTAGAATGC | same as above, but for the mutation V387A |
| PAll1338_V387A_r | GACTTTAATAGCGCGATCGCGCTTGGTAG | |
| PAll1338_I388A_f | TGGCCAAAGTCCCCGATAGTAGAATGCT | same as above, but for the mutation I388A |
| PAll1338_I388A_r | GGGACTTTGGCCACGCGATCGCGCTTG | |
| PAll1338_K389A_f | ATTGCGGTCCCCGATAGTAGAATGCTG | same as above, but for the mutation K389A |
| PAll1338_K389A_r | CGGGGACCGCAATCACGCGATCGCGCT | |

| | | |
|---|---|---|
| PAll1338_V390A_f | TTAAAGCCCCCGATAGTAGAATGCTGCAC | same as above, but for the mutation V390A |
| PAll1338_V390A_r | TCGGGGGCTTTAATCACGCGATCGCGCT | |
| PAll1338_M395A_f | GATAGTAGAGCGCTGCACAAAGCTCGCACC | same as above, but for the mutation M395A |
| PAll1338_M395A_r | GCGCTCTACTATCGGGGACTTTAATCACG | |
| PAll1338_L396A_f | AGTAGAATGGCTCACAAAGCTCGCACCCAT | same as above, but for the mutation L396A |
| PAll1338_L396A_r | GAGCCATTCTACTATCGGGGACTTTAATCA | |
| PAll1338_L403A_f | CATGCCCAAGCTAAGGGTATCACCAGAC | same as above, but for the mutation L403A |
| PAll1338_L403A_r | TAGCTTGGGCATGGGTGCGAGCTTTGTG | |

**Supplementary References**


Battesti A., and Bouveret E. (2012) The bacterial two-hybrid system based on adenylate cyclase reconstitution in *Escherichia coli*. *Methods* **58:** 325-334.

Elhai J., Vepritskiy A., Muro-Pastor A.M., Flores E., and Wolk C.P. (1997) Reduction of conjugal transfer efficiency by three restriction activities of *Anabaena* sp. strain PCC 7120. *J Bacteriol* **179:** 1998-2005.

Felsenstein J. (1985) CONFIDENCE LIMITS ON PHYLOGENIES: AN APPROACH USING THE BOOTSTRAP. *Evolution* **39:** 783-791.

Kumar S., Stecher G., Li M., Knyaz C., and Tamura K. (2018) MEGA x: molecular evolutionary genetics analysis across computing platforms. *Mol Biol Evol* **35:** 1547-1549.

Mitschke J., Vioque A., Haas F., Hess W.R., and Muro-Pastor A.M. (2011) Dynamics of transcriptional start site selection during nitrogen stress-induced cell differentiation in *Anabaena* sp. PCC7120. *Proc Natl Acad Sci U S A* **108:** 20130-20135.

Niu T.C., Lin G.M., Xie L.R., Wang Z.Q., Xing W.Y., Zhang J.Y., and Zhang C.C. (2019) Expanding the potential of CRISPR-Cpf1-Based genome editing technology in the cyanobacterium *Anabaena* PCC 7120. *ACS Synth Biol* **8:** 170-180.

Saitou N., and Nei M. (1987) The neighbor-joining method: a new method for reconstructing phylogenetic trees. *Mol Biol Evol* **4:** 406-425.

Shih P.M., Wu D., Latifi A., Axen S.D., Fewer D.P., Talla E., Calteau A., Cai F., Tandeau de Marsac N., Rippka R., et al. (2013) Improving the coverage of the cyanobacterial phylum using diversity-driven genome sequencing. *Proc Natl Acad Sci U S A* **110:** 1053-1058.

Zhang J.Y., Deng X.M., Li F.P., Wang L., Huang Q.Y., Zhang C.C., and Chen W.L. (2014) RNase E forms a complex with polynucleotide phosphorylase in cyanobacteria via a cyanobacterial-specific nonapeptide in the noncatalytic region. *RNA* **20:** 568-579.

Zhou C., Zhang J., Hu X., Li C., Wang L., Huang Q., and Chen W. (2020) RNase II binds to RNase E and modulates its endoribonucleolytic activity in the cyanobacterium *Anabaena* PCC 7120. *Nucleic Acids Res* **48:** 3922-3934.
